# Gene flow between two thick-billed grasswren subspecies with low dispersal creates a genomic pattern of isolation-by-distance

**DOI:** 10.1101/2021.09.16.460701

**Authors:** Amy L. Slender, Marina Louter, Steven A. Myers, Tessa Bradford, Michael G. Gardner, Sonia Kleindorfer

## Abstract

**Context:** In the era of the Anthropocene, habitat loss and environmental change threaten the persistence of many species. Genotyping-By-Sequencing (GBS) is a useful molecular tool for understanding how patterns of gene flow are associated with contemporary habitat distributions that may be affected by environmental change. Two parapatric subspecies of the threatened thick-billed grasswren (TBGW; *Amytornis modestus*) more frequently occur in different plant communities. As such, a preference for plant community type could reduce subspecific introgression and increase genetic diversity at the parapatric boundary.

**Aims:** We aimed to measure gene flow within and among two TBGW subspecies and tested whether divergent genomic markers were associated with plant community type.

**Methods:** We sequenced 118 individuals from either of the two TBGW subspecies or in the region of parapatry and identified 7583 SNPs through ddRADseq.

**Key results:** We found evidence of asymmetric gene flow and a genomic pattern of isolation-by-distance. There were sixteen genomic outliers correlated with plant community type (regardless of location).

**Conclusions:** These findings show that plant community type does not prevent introgression in one subspecies (*A. m. raglessi*), but low dispersal and habitat heterogeneity could contribute to the maintenance of distinct subspecific morphotypes. Local adaptation in different plant community types could also provide a mechanism for future divergence.

**Implications:** We suggest subspecific introgression could increase genetic variation and the adaptive potential of the species, facilitating species persistence under conditions of climate change.

**Introgression between grasswren subspecies:** Characterising gene flow facilitates conservation management. This study used genomic markers to measure gene flow between thick-billed grasswren subspecies and found results that support taxonomic identification of the two subspecies and suggests grasswrens have low dispersal and may benefit from increased genetic diversity. Recognition of models of divergence with gene flow will be necessary for future conservation management.

## Introduction

Habitat loss is the leading cause of reduced species persistence and species extinction (Bradshaw 2012; Newbold *et al*. 2015; Allan *et al*. 2019; Thompson *et al*. 2019). Within Australia, habitat loss has been anthropogenically driven by a multitude of processes that has changed the landscape notably since the late 18^th^ century. These processes include the introduction of invasive species, anthropogenic dispersal of non-local species, redirection/removal of natural water courses, and changes in soil properties due to agricultural practices (Kingsford 2000; Woinarski *et al*. 2015; Jellinek *et al*. 2020; Mallen-Cooper and Zampatti 2020). An alarming proportion of extant species are threatened by habitat loss, and, consequently, have reduced population sizes and limited genetic variation on which selection can act (Saccheri *et al*. 1998; Amos *et al*. 2012). Molecular tools are important for conservation management practices and species interventions, as they mediate threats to wildlife and ensure long-term success of intervention programs (Elshire *et al*. 2011; Steiner *et al*. 2013; Flockhart *et al*. 2015; Deiner *et al*. 2017; Forseth *et al*. 2017). Population genetics can identify populations that may be in greater need of intervention or better suited for conservation management (Dudgeon *et al*. 2012; Paparella *et al*. 2015; Whiteley *et al*. 2015; Willoughby *et al*. 2015; Rosauer *et al*. 2018; Mynhardt *et al*.2020; Rossetto *et al*. 2021). Understanding how species respond to habitat changes is relevant for mitigating future threats, especially where further habitat change is predicted to occur.

Populations may be more likely to cope with climate change if they are able to expand their range and move into novel habitats (Hoffman and Blows 1994). There are several evolutionary dynamics that determine whether a species can expand their range or not. These include how much genetic variation there is at the population margin, the strength of genetic swamping of genotypes from central to marginal individuals, and the heritability of adaptive traits at the population margin (Jenkins and Hoffman 1999; Davis *et al*. 2013; Moerman *et al*. 2020). Local adaptation into novel environments at the species boundary is one factor that promotes range-expansions, as observed in the European damselfly (*Ischnura elegans*) (Dudaniec *et al*. 2018). Gene flow can erode local adaption that may favour range expansion, but – if the population is large enough – gene flow could also facilitate local adaptation by enhancing genetic variation (Kirkpatrick and Barton 1997; Case and Taper 2000). At the leading margin of the European lizard (*Zootoca vivipara louislantzi*), low gene flow has facilitated a range expansion but low genetic diversity throughout the population could also mean this lizard is susceptible to decline in the face of future climate change (Dupoué *et al*. 2020). When range-shifts involve secondary contact between divergent taxa, species persistence could also be affected due to loss of locally adaptive traits, hybrid swarms or interspecific competition (Case and Taper 2000; Sanchez-Guillen *et al*. 2016). Conservation of threatened species under future ecological scenarios will depend on the ability to predict range shifts, and an understanding of the genomics of hybridisation and introgression.

Associations between populations and their habitat develop through ecological opportunity (Wellborn and Langerhans 2015). For example, morphotypes that give a population an advantage in their particular habitat type are likely to be retained (Aiello *et al*. 2021; Grismer 2021). The strength of an ecological association will be influenced by the amount of gene flow occurring between populations with different ecological associations, which in turn is dependent on ease of dispersal across the landscape. Individuals are more likely to disperse to habitats that are similar to their habitat of origin. This is because individuals that are locally adapted will have lower fitness outside their original habitat type (Fedorka *et al*. 2012; Berner and Thibert-Plante 2015). Therefore, populations occurring in linear, unfragmented landscape arrangements, such as habitat gradients, could have reduced gene flow and in turn stronger ecological associations (e.g. Cicero 2004). Populations that occur in landscapes with more diverse patterns of habitat distribution, such as patchy and heterogeneous landscapes, could have greater gene flow because individuals need to disperse greater distances to reach particular habitat types and could therefore choose to remain in an alternate habitat type (Lenormand 2002; Harrisson *et al*. 2012; Forester *et al*. 2016). It may be less likely for associations between populations and their habitat to occur in a heterogeneous landscape because gene flow will reduce the frequency of locally selected alleles. More case studies are needed to complement a growing body of theoretical modelling, to inform our understanding of the occurrence of ecological associations and the magnitude of gene flow across different landscape scenarios, ultimately with a view to better manage extant populations.

The endangered thick-billed grasswren (*Amytornis modestus,* TBGW) is an arid-zone species of the Maluridae family. We adopt the nomenclature of (Black 2011; 2016) which describes seven subspecies of TBGW. There are two extinct and five extant subspecies occurring in parts of the Northern Territory, South Australia and New South Wales (Black *et al*. 2011; Black and Gower 2017). This taxonomy is a widely accepted (Skroblin and Murphy 2013; Gill and Donsker 2017) however competing taxonomic assignments have been proposed (Christidis *et al*. 2013; Norman and Christidis 2016). Studies show that *A. m. indulkanna* and *A. m. raglessi* are distinct based on morphology and mitochondrial sequences (Austin *et al*. 2013). These subspecies share a region of parapatry between the salt lakes, Lake Eyre and Lake Torrens that likely formed due to secondary contact and a possible range expansion (Slender *et al*. 2017). Outside the region of parapatry, the habitat that each subspecies occupies is characterized by a different and distinct plant community (Slender *et al*. 2018a). Within the region of parapatry, there is a third ‘sandy’ habitat type where grasswrens were rarely present (Slender *et al*. 2018a). The Central Australian arid zone is known for its heterogeneous distribution of different plant types (Slatyer 1961; Williams 1982; Brandle 1998). This feature, along with the habitat changes associated with grazing in the arid zone (Jessop 1995; Navarro *et al*. 2006; Facelli and Springbett 2009), is likely to impact gene flow between populations associated with particular plant communities. In general, the arid zone is predicted to experience greater temperature extremes, less precipitation, and more extreme weather events in the future (Pickup 1998; Lioubimtseva 2004; Lindenmayer and Burgman 2005; Vaghefi *et al*. 2019). Adaptability through greater genetic diversity will be critical for the persistence of the two parapatric TBGW subspecies.

In this study, we aimed to measure gene flow within and among two TBGW subspecies that have been observed in different plant communities (*A. m. indulkanna* in plant community A, dominated by *Maireana aphylla* [cotton saltbush], and *A. m. raglessi* in plant community B, dominated by *M. astrotricha* [low bluebush] and *M. pyramidata* [blackbush]) (Slender *et al*. 2018a). The two subspecies may overlap in an area where a third plant community (plant community AB, dominated by *Zygochloa paradoxa* [sandhill canegrass]) occurs but which is not considered suitable foraging habitat for TBGW (Black *et al*. 2011; Slender *et al*. 2018a). This area, the parapatric margin, has been proposed as an area of secondary contact. We examine whether strength of gene flow changes across the three regions that historically were likely to have been demographically different and today contain different plant community types. We test the idea that gene flow is contemporarily higher in the parapatric margin.

## Materials and Methods

### Samples

We used DNA from all available TGBW samples which included a combination of 104 contemporary samples and 14 museum samples (Table S1; supplemental material). Contemporary samples were collected in the field by mist-netting birds during the breeding seasons from 2012 to 2015. For further details on the study species and contemporary sample collection methods see Slender *et al*. (2017). Museum samples were collected from two time periods; four museum samples were from 1985 (*A. m. raglessi*) and the remainder were from 2007 to 2009 (*A. m. raglessi* [*n* = 2] and *A. m. indulkanna* [*n* = 8]) (Austin *et al*. 2013). Samples were organized into three geographically associated zones described in Slender *et al*. (2017) in order to compare genetic diversity and gene flow between the subspecies centre’s and their parapatric margin (Figure 1). Zone AB describes the subspecies parapatric margin; zone A describes the geographic centre of *A. m. indulkanna* and occurs to the west of zone AB and zone B describes the geographic centre of *A. m. raglessi* and occurs to the east of zone AB. TBGWs in zone A were predominantly found in habitat containing *Maireana aphylla* (cotton bush) and *Atriplex nummularia omissa* (Oodnadatta saltbush) (Black *et al*. 2011; Slender *et al*. 2018a).While TBGWs in zone B were predominantly found in habitat with *M. astrotricha* (low bluebush) and *M. pyramidata* (blackbush) (Black *et al*. 2011; Slender *et al*. 2018a). Zone AB contains shrubs typical of TBGW habitat such as *M. astrotricha* (low bluebush) and *A. vesicaria* (bladder saltbush), but this was heterogeneously distributed among stands of *Zygochloa paradoxa* (sandhill canegrass). The boundary between zone A and zone AB has been extended compared to Slender *et al*. (2017) so that two museum samples (SAMA B55668 and SAMA B55667) that were formerly included in zone A, now fall within zone AB. This is because the landscape in this area was more like the habitat of zone AB (Slender *et al*. 2018a).

**Table 1.**
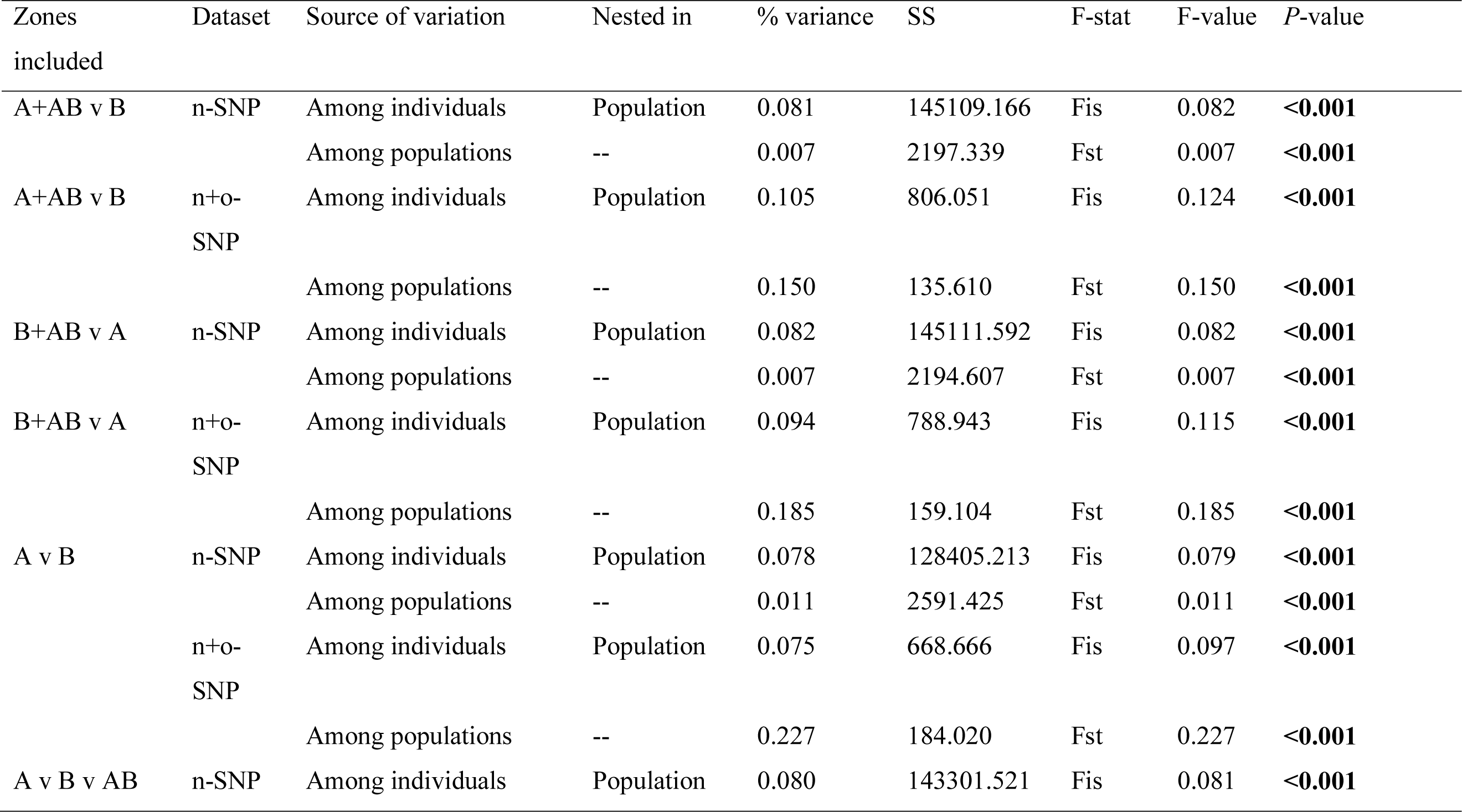
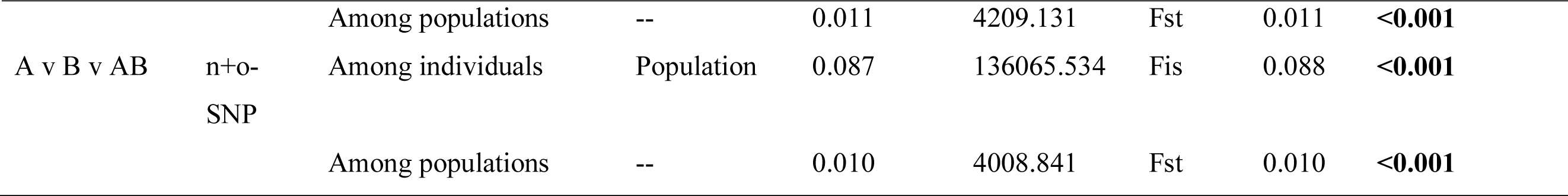
Partitioning of the molecular variance among (1) individuals within zone A and zone B and (2) between zone A and zone B using AMOVA. Zone AB was merged with zone A or zone B in two separate analyses, excluded in a third, or analysed as a separate population. Variance was compared for both n-SNP and o-SNP datasets. Significant *p*-values (< 0.05) are shown in bold.

**Figure 1.**
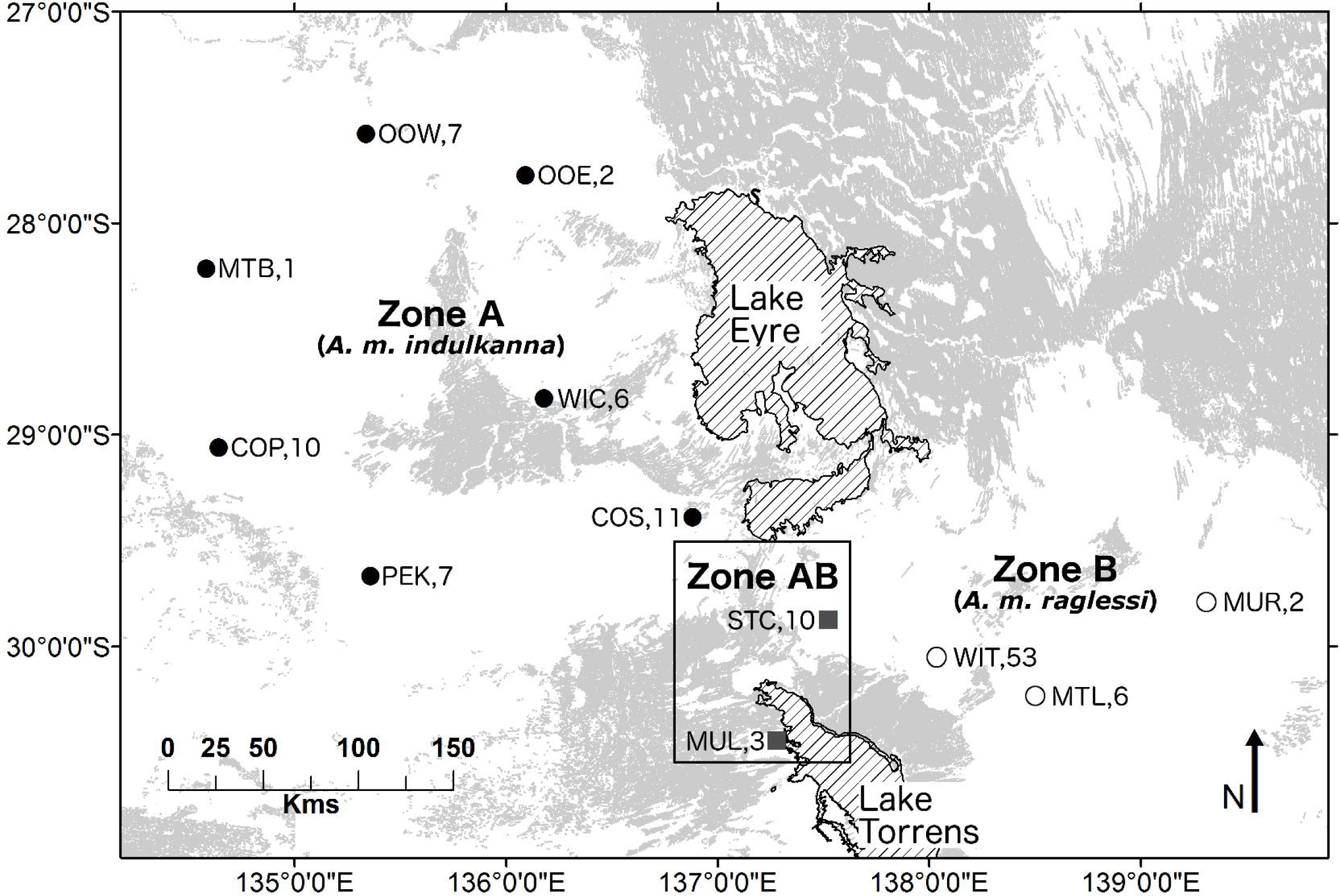
South Australian collection localities for samples of two thick-billed grasswren (TBGW) subspecies used for nuclear genomic sequencing. Localities are grouped into three zones. Sand dunes (grey shade) that run between Lake Eyre and Lake Torrens demarcate a novel habitat type where TBGWs are rarely observed; localities in zone A (solid circle) occur to the west of the sand dunes, localities in zone AB (grey square) occur immediately east of the sand dunes in a region of parapatry (referred to as the contact zone), and localities in zone B occur to the east of zone AB. Locality abbreviations are listed in Table S1. Numbers indicate sample size at that locality.

### DNA extraction

Genomic DNA was extracted from tissue and blood in salt solution using a DNeasy Blood and Tissue kit (QIAGEN Pty Ltd, VIC, Australia) or a Gentra Puregene Blood Kit (QIAGEN Pty Ltd, VIC, Australia). Genomic DNA was extracted from FTA samples following Smith and Burgoyne (2004). DNA extractions were carried out in a separate PCR free laboratory in order to minimise DNA contamination. DNA quantity was measured using the Qubit fluorometer (ThermoFisher Scientific Australia Pty Ltd, VIC, Australia). DNA extractions were quality tested using UV-spectrophotometry and agarose gel electrophoresis. Samples were assessed as good quality when they showed 1) a large un-degraded band on an agarose gel and 2) a 260/280 ratio between 1.8 and 2.0 indicating minimal protein and chemical contamination.

### Library construction and sequencing

Genotyping-by-sequencing libraries were generated following the protocol in Poland *et al*. (2012). DNA samples (200 ng) were digested with 8 U of PstI and MspI at 37°C for 2 hrs. Each sample was prepared for multiplexing by ligating a pair of adapters containing a unique barcode to the DNA fragments. We used 96 unique barcodes where the barcodes ranged from 4 to 9 bp (Elshire *et al*. 2011) to create two pooled libraries. One barcode in each library was assigned as a negative control and seven barcodes in each library were used to duplicate samples within (6 samples) and across (1 sample) libraries. Barcodes were randomly allocated to samples from different geographic locations so that we would detect errors caused by mismatched barcodes that can be made during library preparation or subsequent demultiplexing. We used an adapter mix to DNA ratio of 1:50 ng as this concentration produced libraries with reduced adapter dimer (Elshire *et al*. 2011). Libraries were then amplified using PCR with the following standard Illumina primers: P5 (5’-AATGATACGGCGACCACCGAGATCTACAC-3’) and P7 (5’-CAAGCAGAAGACGGCATACGAGAT-3’). Sequencing was performed on an Illumina next-seq sequencer that produced single end-reads of 62 bp after adapter trimming. Sequencing data was quality checked using FastQC v10.1 (http://www.bioinformatics.babraham.ac.uk/projects/fastqc/).

### SNP calling and filtering

Read filtering and SNP calling was performed using STACKS v1.44 (Catchen *et al*. 2013). Samples were demultiplexed using the *process_radtags* program and reads from sample replicates were merged into one sample (after preliminary SNP calling with separated duplicates was used to determine error rates). Reads were identified if the adapter barcode (with a maximum of 2 mismatches) and the unique barcode (with a maximum of 1 mismatch) were present. Putative alleles were identified from a stack assembly created with the *ustacks* program that was instructed to include loci with a minimum depth of coverage of 5 reads, maximum distance of 2 nucleotides, and maximum number of 50 stacks per locus. The *cstacks* program was used to create a catalog for identifying loci with a maximum of 2 mismatches between putative alleles. SNPs were determined by comparing the output of *ustacks* with the output of *cstacks* using the *sstacks* program. Relaxing the error tolerance rate improves the likelihood of detecting heterozygous calls (Hohenlohe *et al*. 2010; Lu *et al*. 2013). We used a bounded model for detecting SNPs with the lower error limit of 0.0001 and an upper error limit of 0.05. Minor alleles with low frequency cause problems in population genetic analyses because they can represent sequencing error and they are not informative population markers (Gonçalves da Silva *et al*. 2015). We removed loci (1) that were missing calls in more than 80% of all individuals, or (2) if the minor allele frequency was < 0.05. An individual was considered heterozygous at a locus if there was a proportion of <0.75 reads per allele. We checked that the dyadic likelihood of relatedness did not exceed 0.4 between any individual within zone A and zone AB or zone B and zone AB using the program COANCESTRY v1.0.1.2 (Wang 2011). A related individual of a pair or group of related individuals was excluded if they were related to more individuals and if they had more missing data.

The output from STACKS consisted of 16,569 loci that we applied additional filtering steps to with a custom script implemented in R STUDIO v1.0.136 (R Core Development Team 2008). Loci were removed if they appeared in the negative control and were observed in less than 85% of samples. We used a Principal Component Analysis (PCA) in the R package *adegenet* v2.0.1 (Jombart 2008) to explore preliminary population structure. The putative clusters without admixed individuals were each analysed for loci out of Hardy-Weinberg Equilibrium (HWE) in the R package *pegas* v0.9 (Paradis 2010). We removed loci from further analysis that did not conform to HWE in (1) both putative clusters or (2) one putative cluster when a SNP was only present in one cluster. We identified linked loci in each putative cluster excluding potentially admixed individuals, using PLINK v1.07 (Purcell *et al*. 2007). We removed loci from further analysis that were highly correlated (r^2^ > 0.1) and had a p-value <0.01 in (1) both putative clusters or (2) one putative cluster when a SNP was only present in one cluster. Within a linkage pair, we removed the locus with the most linkage pairs. When both loci had even numbers of linkage pairs, we removed the locus with the most missing data.

### Differences between putative genetic clusters

*F_ST_* outlier loci between putatively non-admixed individuals in zone A and zone B were identified using two programs. We ran BAYESCAN v2.1 (Foll and Gaggiotti 2008) with default settings after data format conversion with PGDSPIDER v2.1.1.0 (Lischer and Excoffier 2012) and the R package *OutFLANK* v0.1 (Whitlock and Lotterhos 2015). *F_ST_* outlier loci were defined as having a *q*-value and corresponding false discovery rate of < 0.1. Using a consensus list of *F_ST_* outlier loci from both analyses, the dataset was separated into three versions, one with neutral loci (n-SNP), one with only outliers putatively under selection (o-SNP), and a third with both neutral and outlier loci (n+o-SNP). The closest known species relative with an available whole genome sequence is the zebra finch (*Taeniopygia guttata*) (Warren *et al*. 2010). We performed a discontiguous megablast search that looked for sequence similarities between TBGW o-SNPs and the zebra finch GenBank and refseq assemblies using the blastn and blastx functions respectively with an evalue threshold of 1e-6.

To further understand the distribution of shared and distinct genetic variation, we performed an Analysis of Molecular Variance (AMOVA) and calculated the significance of pairwise *F*_ST_ between zones using GENODIVE v2.0b27 (Meirmans and Van Tienderen 2004) with 10,000 permutations. We tested differences between genetic clusters in three separate analyses; one where the region of parapatry was merged with zone A, one where the region of parapatry was merged with zone B and the last where zone AB was excluded. We repeated these analyses with the n-SNP dataset and n+o-SNP dataset. Expected heterozygosity (H_e_) is a measure of genomic diversity when the dataset consists of SNPs (Fischer *et al*. 2017). H_e_ was calculated for each zone separately using the n+o-SNP dataset.

### Isolation-By-Distance

We tested for Isolation-By-Distance (IBD) among eleven sampling localities by calculating geographic and genetic distance matrices that excluded the locality MTB (zone A) as it contained only one individual (Figure 1). The Euclidean distance between localities (km) was first calculated in GENALEX v6.5 (Peakall and Smouse 2006; Peakall and Smouse 2012). Any paths between localities that passed through Lake Eyre or Lake Torrens (e.g., MUL and WIT) were corrected so that it did not pass through the salt lake. This was done by calculating the Euclidean distance from the first sampling location to a point in the middle of the space between Lake Eyre and Lake Torrens and then calculating the Euclidean distance between that point and the second sampling location and adding the distances together. All geographic distances between sampling localities were then log transformed to account for individuals moving in two dimensions. We calculated a pairwise *F*_ST_ genetic distance matrix with n-SNPs using GENODIVE (Meirmans 2020) and also transformed the genetic data (*F*_ST_/1 – *F*_ST_) (Nei 1977). Tests for IBD are easily biased by hierarchical population structure where allele frequencies are sharply divided geographically (Meirmans 2012) as well as uneven sample sizes and the spatial patterns between sampling localities (Balkenhol *et al*. 2009; Guillot and Rousset 2013; Kierepka and Latch 2015). We therefore performed a series of tests for IBD using three different methods; (1) Mantel and partial Mantel tests, (2) Decomposed Pairwise Regression (DPR), and (3) spatial autocorrelation. To test for limitations in gene flow that might prevent genetic swamping at marginal locations we performed two Mantel tests: (1) across locations within zone A (*A. m. indulkanna*) and zone AB, and (2) across locations within zone B (*A. m. raglessi*) and zone AB. We included zone AB in an analysis with either zone because this area appears to be the population margin for both subspecies (the region of parapatry) (Slender *et al*. 2017). We then performed a partial Mantel test across all zones to test for gene flow among subspecies while accounting for potential population structure across these regions. We used a binary matrix that compared zone B versus zone AB and A combined or zone A versus zone AB and B combined. We used GENODIVE (Meirmans 2020) to perform mantel and partial mantel tests with 1000 permutations. DPR is useful for detecting outlier populations that may be associated with weak geographic barriers such as heterogeneous landscapes (Koizumi *et al*. 2006). We performed a DPR using the R package *DPR* v1.0 (Reynolds 2011).

Finally, we used spatial autocorrelation (Smouse and Peakall 1999) in GenAlEx v6.5 to further evaluate spatial structure in the genetic data at an individual level. A pairwise matrix with Roussets’s *a* genetic distance (Rousset 1997; Rousset 2000) between all individuals with the n-SNP dataset was calculated using SPAGeDi v1.4b (Hardy and Vekemans 2002). Geographic distances between individuals were calculated in GenAlEx using the same method to create the geographic distance matrix for the mantel tests. Distance classes were sufficiently small enough to evaluate any non-linear correlations with the autocorrelation coefficient (*r*) where the sample size within each distance class was relatively even. We looked for the presence of IBD within each distance class as well as the detectability of IBD across multiple distance classes (Diniz-Filho and Pires de Campos Telles 2002). Significance was assessed for both tests using 95% confidence intervals for the null hypothesis of no spatial structure using 999 random permutations, and for estimates of *r* by bootstrapping 999 pairwise comparisons for each distance class.

### Gene flow

We investigated population structure and admixture using the n-SNP dataset with two methods: (1) Discriminant Analysis of Principal Components (DAPC) (Jombart *et al*. 2010) implemented in the R package *adegenet* v2.0.1 (Jombart 2008), which assigns individuals to genetic clusters following a PCA while accounting for within-population variation; and (2) Bayesian clustering with the program STRUCTURE v2.3.4 (Pritchard *et al*. 2000; Falush *et al*. 2003) that determines genetic clustering based on HWE. All methods are useful for detecting admixed individuals. For the DAPC, we retained one principal component, as this returned the optimum *a*-score, which is the difference between the proportion of successfully reassigned individuals compared to the number of principal components retained. The optimum number for *K* was inferred from the retained principal component by identifying *K* where the Bayesian Information Criterion (BIC) produced an elbow in the curve of BIC values as a function of *K*. Admixture was inferred if the proportion of population assignment was <0.9 or >0.1 in any individual. For the STRUCTURE analysis, three replicate runs for each *K* were analysed (as Standard Deviation of LnP(K) was small) using default settings, unless stated. We used the admixture model with correlated allele frequencies and an MCMC chain of 1,000,000 iterations with a burnin of 10,000 iterations to test *K* between 1 and 5. To estimate the probability of mixed ancestry for each individual, the option ANCESTDIST was used. Admixture was inferred if the confidence intervals of the individual population assignment did not include 1 or 0 in all three replicate runs. STRUCTURE HARVESTER (Earl and vonHoldt 2011) was used to estimate the best fitting value for *K*. When the highest LnP(*K*) was not *K* = 1, then the most likely *K* was determined using Delta *K* (Evanno *et al*. 2005). Cluster assignments were merged in CLUMPP v1.1.2 (Jakobsson and Rosenberg 2007) and results were visualized with DISTRUCT v1.1 (Rosenberg 2004). Population assignments of individuals were compared to their mtDNA hapolotype (Slender *et al*. 2017). Further hierarchical population structure was investigated by repeating the analysis on individual populations detected in the initial run (Evanno *et al*. 2005).

### Selection

Previous comparisons of habitat within the three zones identified three predominant plant communities represented by three principal components (Slender *et al*. 2018a). We used Latent Factor Mixed Models (LFMMs) (Frichot *et al*. 2013) to test for associations between genotype (n+o-SNPs) and the environmental variables defined by the principal components. The LFMM test was performed in the R package *lfmm* v0.0.

### Migration

We tested the proportion of migrants between the three zones with a reduced dataset of 200 loci in BAYESASS v3.0.4 (Wilson and Rannala 2003). We performed a PCA with all individuals in the n-SNP dataset with the R package *adegenet* v2.0.1 (Jombart 2008) and selected loci for use that had the highest loading in the PCA. Three independent MCMC runs were performed with 1,000,000 iterations and a burn-in of 10,000 iterations. The alpha (allele frequency) and delta (inbreeding coefficient) values were adjusted to 0.6 and 0.4 respectively so that the acceptance rates were between 20% and 60%. Iterations were sampled every 100 intervals to determine the posterior distribution of the parameters. Convergence of the MCMC run was assessed by inspecting the trace file in the program TRACER v1.6.0 (Rambaut *et al*. 2015).

## Results

### DNA extraction and sequencing statistics

We used samples from across three zones: zone A (*n* = 44), zone B (*n* = 61), and zone AB (*n* = 13) to assess gene flow between the two parapatric TBGW subspecies. Following DNA extractions, samples stored on FTA® cards produced considerably lower quantities of DNA (<500 ng) compared to blood stored in salt solution (>1,000 ng). Following illumina sequencing, the average number of reads per sample (before filtering) was 2,539,005 with a coefficient of variation of 24.6%. The average between run reproducibility, calculated by determining when the genotype was the same in duplicates on different plates, was 95.7% (*n* = 12,192 loci, excluding missing genotypes). The average within run reproducibility, calculated by determining when the genotype was the same in duplicates on the same plate, was 90.5% (*n* = 146,304 loci, excluding missing genotypes). The average genotyping error rate, calculated from the number of allelic mismatches across duplicates, was 0.31% (*n* = 316,992 loci). There were no individuals that exceeded > 30% missing data; overall the dataset contained 5.56% missing data. Following SNP calling in STACKS, we removed 5 loci that appeared in the negative control and 2929 loci that had low coverage across samples. An initial PCA showed two putative genetic clusters with individuals from zone A forming one cluster, individuals from zone B forming the second cluster and 19 potentially admixed individuals from zone AB and zone B (Figure S1). We observed similar amounts of missing data between the clusters, excluding potentially admixed individuals (cluster 1 [zone A]: 6.41%, cluster 2 [zone B]: 6.58%). We removed a further 625 loci from further analysis that did not conform to HWE and 5428 loci that could potentially introduce linkage disequilibrium.

### Subspecies variation

The proportion of heterozygous SNPs per sample varied from 0.234 in a sample from zone A to 0.301 in samples from both zone A and zone B. Mean ± SE estimates of heterozygosity (He) were slightly higher for zone A and zone B compared to zone AB (zone A = 0.303 ± 0.002, zone B = 0.304 ± 0.001; zone AB = 0.288 ± 0.002). The number of private alleles within zone B was greater (n = 16) than for either zone A (n = 1) or zone AB (n = 0). We identified 39 loci as potential *F_ST_* outliers under selection which left 7543 loci that were treated as neutral loci not under selection. Therefore, the dataset o-SNP contained 39 loci and the dataset n-SNP contained 7543 loci. Of the 39 outlier loci, nine loci were monomorphic in zone A; three loci were monomorphic in zone B and four were monomorphic in zone AB. Of the four monomorphic loci in zone AB, three were shared with the monomorphic outliers of zone A and one was shared with the monomorphic outliers of zone B. Four outliers had hits to nucleotide sequences from the zebra finch GenBank assembly (Table S2) but there were no matches to protein sequences from the refseq assembly. These blast hits did not reveal why there could be associations between outlier loci and plant community type. Zone B had slightly more polymorphic loci in the n-SNP dataset (99.9%) compared to zone A (99.2%). Using the n-SNP dataset, the proportion of total genetic variance was shared among individuals and populations similarly when the region of parapatry (zone AB) was combined with either zone A or zone B, or even when it was excluded (Table 1). The proportion of variance in the case of n-SNP was greater among individuals (mean 0.080%, *p* < 0.001) than among populations ( mean 0.008%, *p* < 0.001; (Table 1). Using the n+o-SNP dataset, the proportion of total genetic variance explained by population was greater than that explained among individuals for all three tests (A+AB v B, B+AB v A, A v B). This difference was greatest when zone AB was excluded. When zone AB was not excluded the difference was greater when combined with zone B (among individuals = 0.185%, *p* < 0.001; among individuals = 0.094%, *p* < 0.001). Using the n-SNP dataset, there was no difference in pairwise estimates of *F*_ST_ when the region of parapatry was combined with either zone A or zone B (Table 2). The pairwise estimates of *F*_ST_ using n+o-SNP was higher when zone AB was combined with zone B (0.202, *p* < 0.001) compared to when zone AB was combined with zone A (0.165, *p* < 0.001; Table 2).

**Table 2.** Pairwise differentiation when zone AB is merged with zone A, zone AB is merged with zone B or zone AB is excluded for both the n-SNP and n+o-SNP datasets. Cells show *F*_ST_ and *p*-values in parentheses. *P*-values were calculated after 10,000 permutations. Significant *p*-values (< 0.05) are shown in bold.

| Zone | n-SNP | n-SNP | n+o-SNP | n+o-SNP |
| --- | --- | --- | --- | --- |
|  | A | B | A | B |
| B+AB | 0.008 (**<0.001**) | -- | 0.202 (**<0.001**) | -- |
| A+AB | -- | 0.008 (**<0.001**) | -- | 0.165 (**<0.001**) |
| A | -- | 0.010 (**<0.001**) | -- | 0.227 (**<0.001**) |

### Isolation-By-Distance

IBD was detected in only one Mantel test that included localities from zone A and zone AB (R^2^ = 0.112, Rxy = 0.335, *p* = 0.029) (Figure S2). There was no correlation between genetic and geographic distance across localities from zone B and zone AB (R^2^ = 0.012, Rxy = 0.110, *p* = 0.435). However, this result may have been affected by the small number of localities used in this test (Figure S2). Partial Mantel tests across all zones where zone AB was in the same cluster as A or B were significant (zone A+AB vs B; R^2^ = 0.317, Rxy (spearman’s r) = 0.514, *p* = 0.001 and zone B+AB vs A; Rxy = 0.385, *p* = 0.015) (Figure 2). The sample sizes for the spatial autocorrelation were skewed for the lowest distance class (0-20 kms) but for all other distance classes the sample size was on average (± SD) 293 ± 132. This analysis showed that at an individual level there was positive spatial autocorrelation for the first two distance classes (0-20 and 20-40 kms) (Figure 3). When plotting *r* as a function of increasing distance classes, the curve intercepted the *x*-axis at 123.6 kms (Figure 3). IBD was detectible from 0 - 60 km and between 80-100 and 140-160 km (*p* < 0.01). This suggests that spatial autocorrelation is linear up to 60 kms and non-linear at other intervals, which may indicate a pattern of low habitat connectivity. Initial results of the DPR analysis suggested that there were no populations that had greater divergence than what was expected based on distance alone. The model with the smallest AIC_C_ where R^2^ was also the highest and where ΔAIC_C_ < 2 was for 3 sub-populations (OOW, MTL, and MUR) to be potential outliers however this was not significant (Table 3). Regression of all sub-populations with all other sub-populations also suggested genetic drift and gene flow were in equilibrium and no population structure was present.

**Figure 2.**
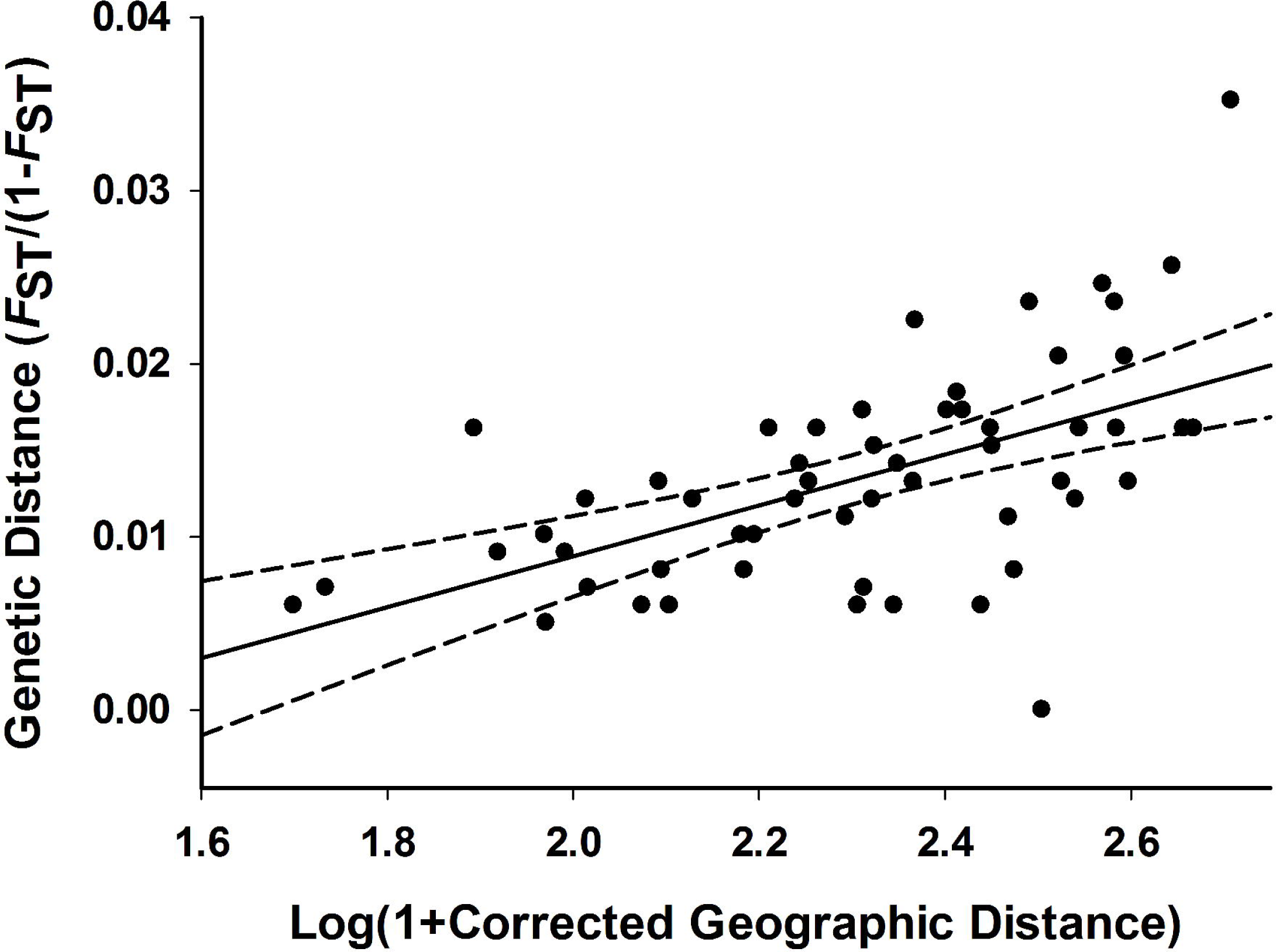
The pairwise genetic (*F*_ST_/(1 – *F*_ST_)) and geographic (log(1 + km)) relationship between localities by zone (zone A: *n* = 6, zone B: *n* = 3, and zone AB: *n* = 2) using a Mantel test (R^2^ = 0.317). There was only one sample collected at the locality MTB, therefore this locality was excluded. The solid line is the line of best fit and the broken lines are the 95% confidence intervals.

**Figure 3.**
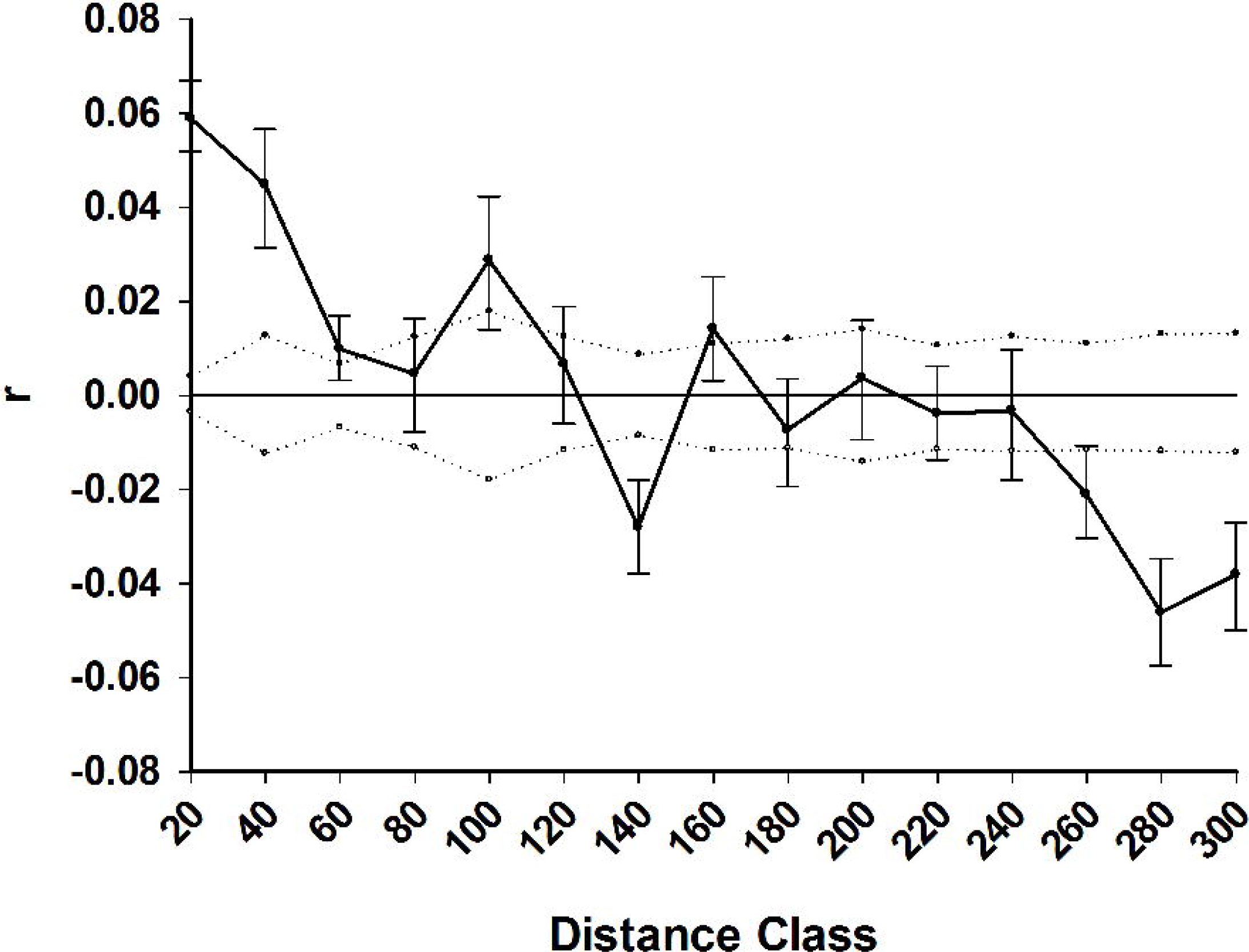
Correlogram showing the spatial autocorrelation coefficient *r* as a function of distance (km) indicated by distance class (end point). Dotted lines are the 95% CI about the null hypothesis of a random distribution of genotypes and error bars are 95% CI of *r*.

**Table 3.** Fit of alternative isolation by distance models with and without putative outlier populations (see Figure 1 for population codes) *n* shows the number of populations in the model, AIC_C_ shows the corrected Akaike’s information criteria, ΔAIC_C_ shows the difference in AIC_C_ between alternative models. These values are used to assess the most likely model. Models are ranked from highest to lowest.

| Population excluded | $n$ | $R^2$ | $P$ | $AIC_C$ | $\Delta AIC_C$ |
| --- | --- | --- | --- | --- | --- |
| MUL, COP, OOE, WIC, WIT, COS, STC, PEK | 3 | 0.92 | 0.182 | -26.635 | 0 |
| MUL, COP, OOE, WIC, WIT, COS, STC | 4 | 0.883 | 0.005 | -39.507 | -12.872 |
| MUL, COP, OOE, WIC, WIT, COS | 5 | 0.825 | <0.001 | -48.503 | -21.868 |
| MUL, COP, OOE, WIC, WIT | 6 | 0.821 | <0.001 | -59.135 | -32.500 |
| MUL, COP, OOE, WIC | 7 | 0.795 | <0.001 | -67.610 | -40.975 |
| MUL, COP OOE | 8 | 0.742 | <0.001 | -76.613 | -49.978 |
| MUL, COP | 9 | 0.575 | <0.001 | -82.782 | -56.148 |
| MUL | 10 | 0.434 | <0.001 | -89.224 | -62.589 |
| None | 11 | 0.317 | <0.001 | -94.404 | -67.770 |

### Geneflow

A PCA showed limited population structure between zone A and B along the first component (1.7% of variation) as there was no separation of individuals into clusters (Figure 4). Despite this, STRUCTURE identified two major genetic clusters (Table S3) corresponding to eastern and western populations. Two genetic clusters were also identified by the DAPC analysis albeit with weaker support (Figure S3). Using *K* = 2, results from both STRUCTURE and the DAPC were concordant in that both analyses showed that 1) zone AB contained the highest proportion of admixed individuals 2) there were greater proportions of admixture in individuals in zone AB than either zone A or zone B, and 3) there were greater proportions of admixture in individuals in zone B than in zone A (Figure 5). Comparison of the two methods showed there were discrepancies in the identity of admixed individuals as well as in the proportions of admixture. The DAPC method compromises the power for detecting admixture with the assignment of individuals to populations, therefore we have limited the discussion of admixture below to the STRUCTURE results. In zone A, 2.3% of individuals were admixed and these individuals had a relatively low proportion of assignment probability from the eastern genetic cluster (< 18%). In zone B, 18% of individuals were admixed and these individuals had low to high proportions of assignment probability from the western genetic cluster (18.7 – 52.0%). Two of the admixed individuals in zone B came from museum samples that were either collected in 1985 or 2007 and were from localities furthest from the region of parapatry (MUR and MTL). In zone AB, all individuals were admixed and had low to high levels of assignment probability from both the eastern (17.5 – 71.4%) and western genetic clusters (28.6 – 82.5%). To look at hierarchical substructure within the identified populations, individuals in zone B and then zone A were excluded from two separate STRUCTURE analyses. For zone A and zone AB, *K* = 1 was the most likely using mean LnP(*K*) and for zone B and zone AB, *K* = 3 was most likely using Delta *K* (Figure S4). Two of smaller clusters from the zone B and zone AB analysis comprised of groups of individuals that were from the same or neighbouring territories and had slightly higher levels of relatedness. An earlier analysis with COANCESTRY showed that the Dyadic likelihood and the 95% confidence intervals for those groups were: *r* = 0.28 (0.26,0.30) – 0.30 (0.28,0.32) for three individuals in the first cluster and *r* = 0.14 (0.12,0.16) – 0.28 (0.25,0.30) for seven individuals in the second cluster. The three individuals in first cluster were also separated along component two (PC2; 1.4% variance) of the PCA (Figure 4).

**Figure 4.**
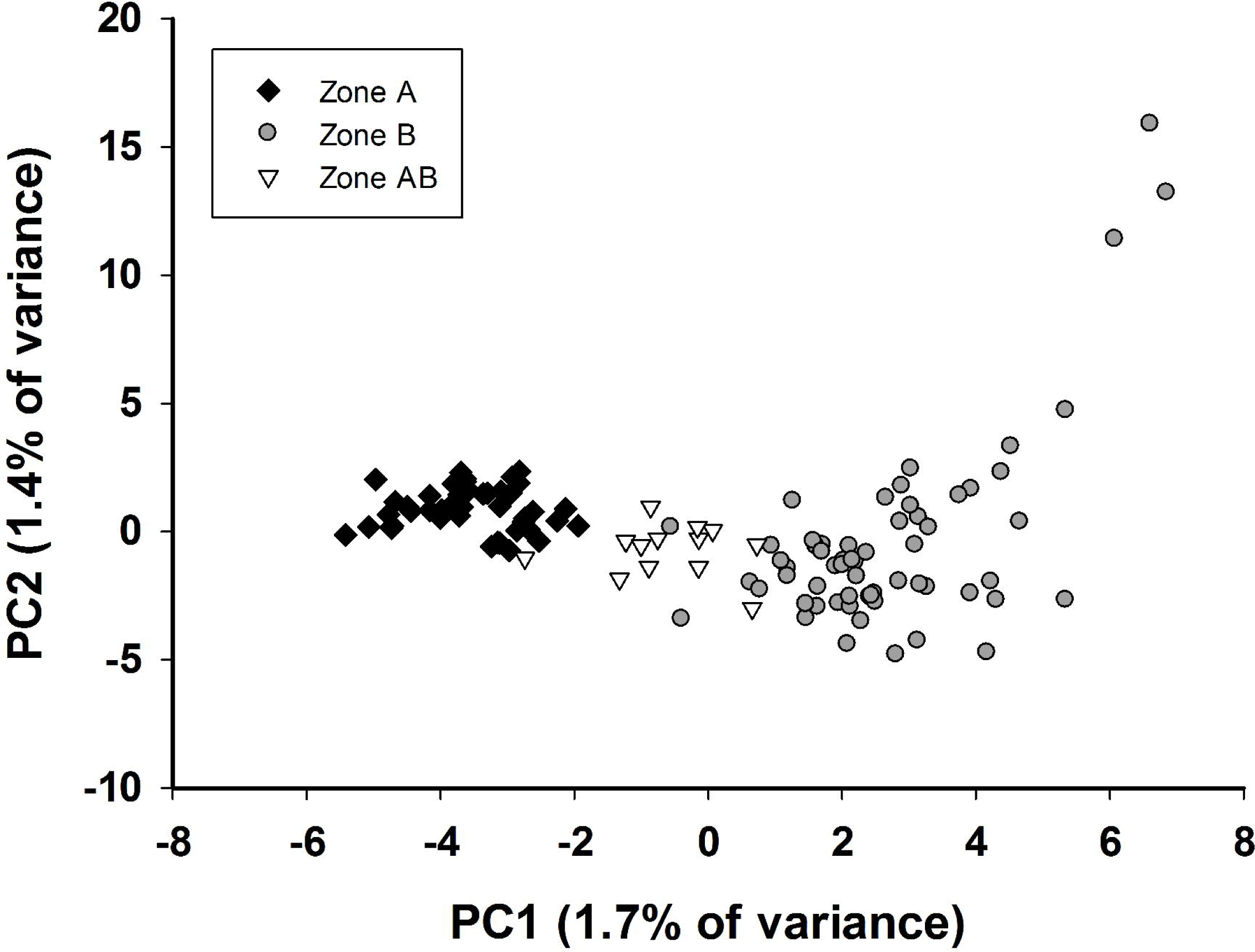
PCA of 7543 loci where individuals from different zones are indicated with different shapes; zone AB (region of parapatry) are white triangles, zone A (*A. m. indulkanna*) are black diamonds, and zone B (*A. m. raglessi*) are grey circles.

**Figure 5.**
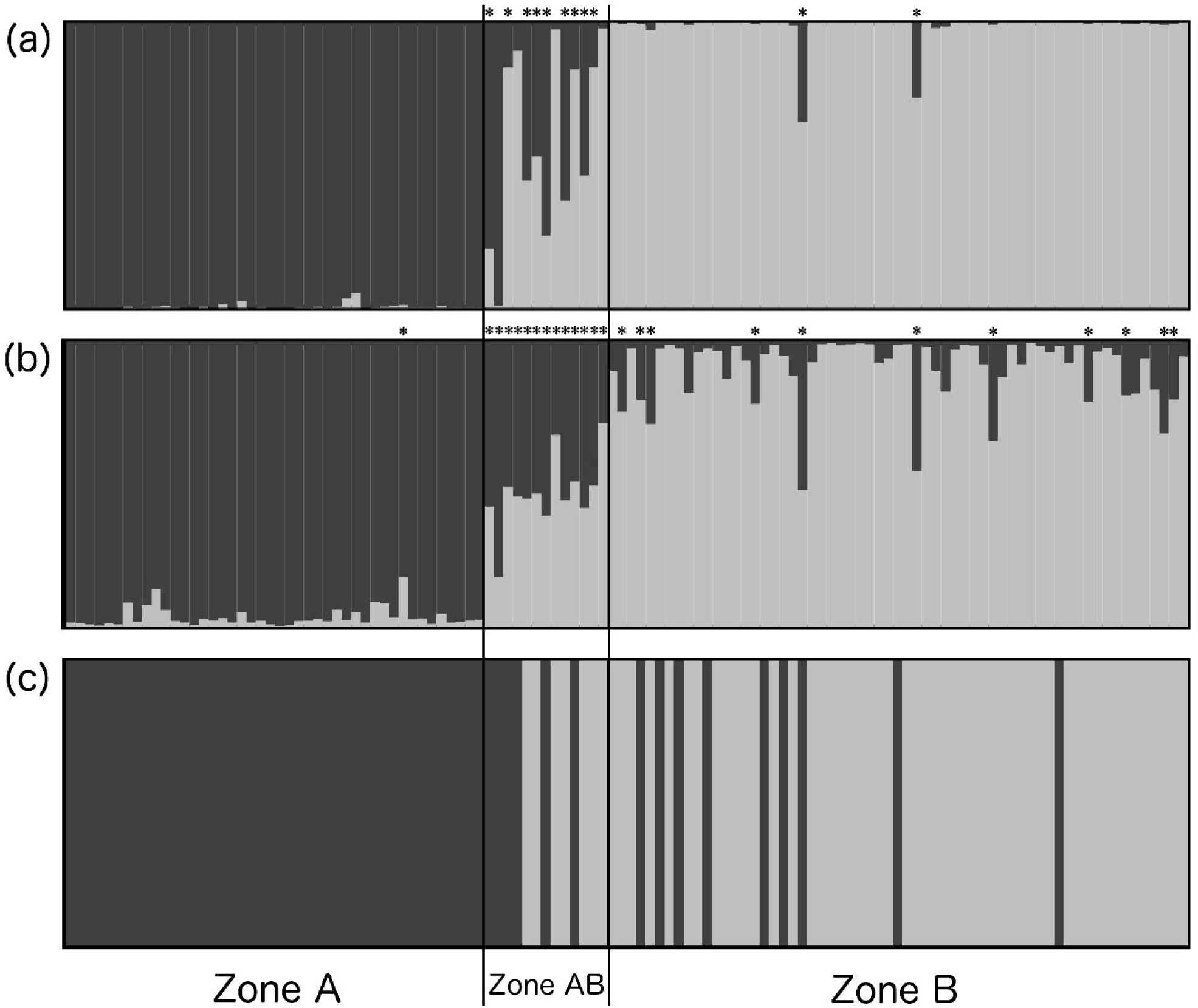
Population assignment tests using 7543 n-SNP loci where *K* = 2 for (a) DAPC and (b) STRUCTURE or using (c) mitochondrial haplotype for ND2 across three zones (Figure 1). Individuals are ordered by latitude in the order listed in Table S1. The proportion of each colour shows the posterior mean proportion of ancestry from the subspecies *A. m. indulkanna* or western haplotype (dark grey) and *A. m. raglessi* or eastern haplotype (light grey). Individuals marked with an asterisk were identified as admixed.

### Ecological associations and migration

A unique plant community was previously identified in each of the three zones using a PCA reported in Slender *et al*. (2018a). PC1 was associated with low abundance of *Atriplex vesicaria* and high abundance of *Zygochloa paradoxa* and was predominant in Zone AB. PC2 was associated with high abundance of *Maireana aphylla* and low abundance of *M. astrotricha* and *M. pyramidata* and was predominant in Zone A. PC3 was associated with low abundance of *A. nummularia omissa* and high abundance of *Acacia* spp and *Rhagodia spinescens* and was predominant in Zone B (Table S4). Using *K* = 2 output from structure, the LFMM analysis identified 328, 333 and 419 loci associated with PC1, PC2 and PC3 respectively. Of the 39 *F*_ST_ outliers, there were 12 loci that correlated with PC2 (two of these also correlated with PC3) and six loci that correlated with PC3. No loci were found to correlate to PC1. The results from BAYESASS suggested that zone AB received more migrants per generation than zone A or zone B (Figure 6). Zone AB received more migrants per generation from zone A than zone B; the mean ± SD migration from zone A = 21.0 ± 4.7% and from zone B = 10.3 ± 4.6%. Zone B received some migration per generation from zone A (4.5 ± 1.6 %), but zone A received < 1% migration per generation from either zone B or zone AB.

**Figure 6.**
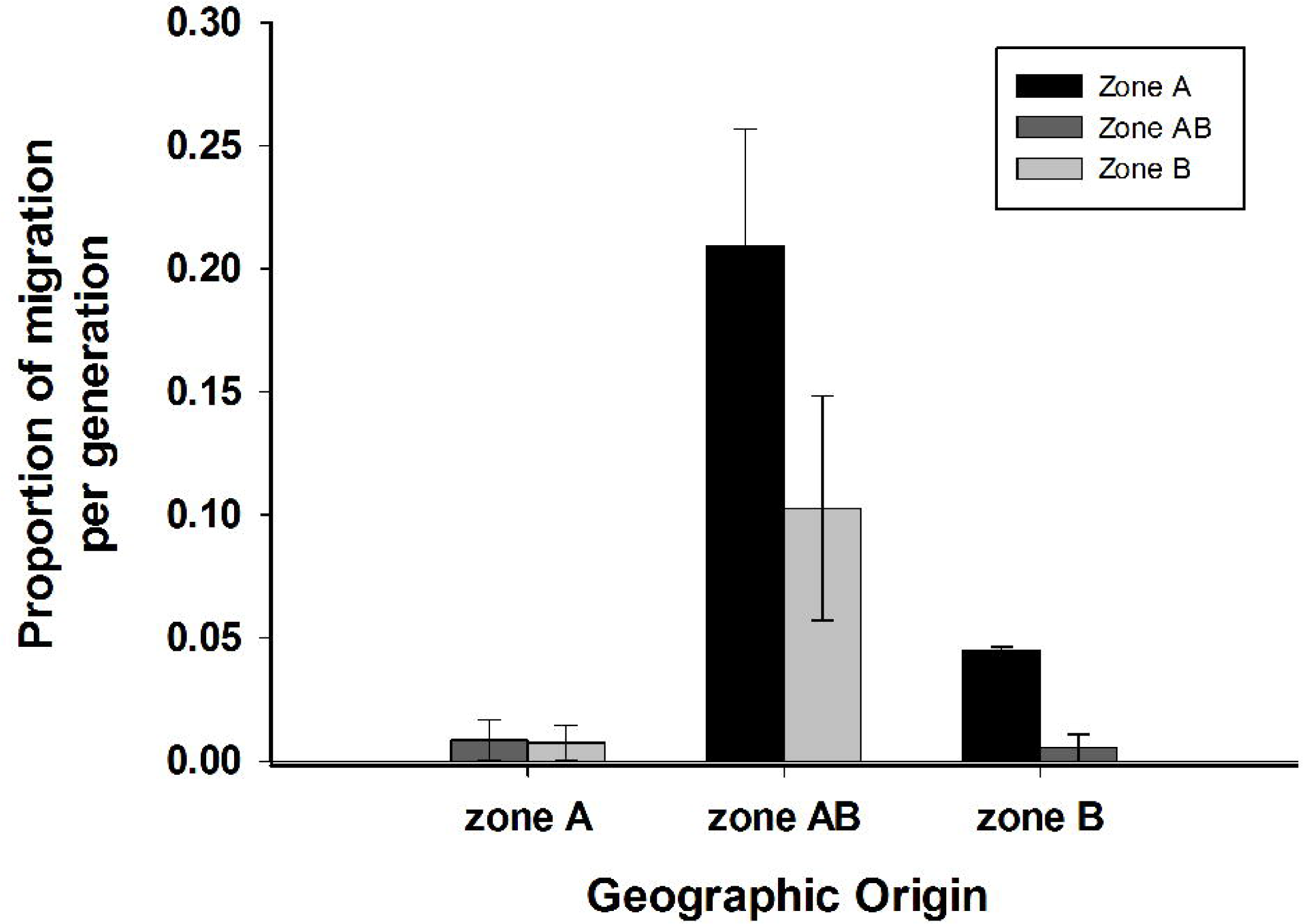
The proportion of migrants (average and standard deviation) assessed between each zone (zone A, zone B, and zone AB) with BAYESASS. Migration from zone A is in black, migration from zone B is in light gray, and migration from zone AB is in dark gray. The analysis was performed following a PCA to identify 200 loci with the highest loading that were used in the BAYESASS analysis.

## Discussion

This study aimed to measure patterns of genetic diversity between the parapatric margin (zone AB) of two TBGW subspecies, and their population centre’s (*A. m. raglessi* = zone B and *A. m. indulkanna* = zone A). Greater genetic variation at the margin could increase the potential for local adaptation to occur in different vegetation types at the margin. We detected gene flow occurring between the subspecies that was not restricted to zone AB as was previously thought and observed no evidence for greater diversity in zone AB compared to other zones. We discovered a pattern of IBD across the subspecies, low genotypic evenness and low genetic differentiation at neutral SNPs based on F_ST_ values indicating the subspecies have introgressed considerably. Spatial autocorrelation at short distances suggests that IBD is likely caused by short-range dispersal. We detected more migration between the parapatric margin and the population centre of *A. m. raglessi* suggesting introgression was asymmetric towards *A. m. raglessi*. There was evidence of local adaptation in both subspecies to different plant communities, which suggests selection could lead to future differentiation of the subspecies.

IBD increases genetic variation because it occurs when there is low gene flow between distant locations. The presence of IBD indicates that individuals within a population only disperse short distances (Aguillon *et al*. 2017). Grasswrens are thought to have poor dispersal ability due to their small size and short wings and have highly localized taxonomies (Christidis *et al*. 2010; Austin *et al*. 2013). This study found evidence for IBD across the TBGW subspecies, *A. m. raglessi* and *A. m. indulkanna*, which have been geographically isolated in the past and have subsequently made secondary contact (Austin *et al*. 2013; Slender *et al*. 2017). The population structure demonstrated in this study is likely biased by the presence of IBD, as limited sampling across large areas replicates patterns of population structure (Perez *et al*. 2018). Poor dispersal is likely to be one mechanism that has created IBD between these subspecies; however, we also detected patterns suggesting landscape heterogeneity could influence gene flow strength. Further work could assess landscape effects on gene flow strength (van Strien *et al*. 2015).

IBD in this study indicates considerable nuclear introgression between the subspecies and a low risk of outbreeding depression (Frankham 2010). Introgression may have ensued over a long period of time if secondary contact between *A. m. indulkanna* and *A. m. raglessi* occurred a long time ago. Alternatively, there may be a preference for heterospecific mates which could also have led to increased introgression. We previously found that *A. m. indulkanna* more often and more intensely responded to hetero-subspecific song than con-subspecific song (Slender *et al*. 2018b). While we know little about the function of grasswren song, it is plausible that greater response to song could indicate mating preferences (Nowicki and Searcy 2005). Introgression of taxonomically young lineages such as subspecies could increase their genetic diversity and the adaptive potential (Grant and Grant 2019). Acknowledging populations that interbreed for conservation planning is a useful component of biodiversity management strategy that is gaining traction in conservation programs (Chan *et al*. 2019). This stands in contrast to previous concerns that introgression is a threat to biodiversity such as when anthropogenic interference creates conditions that promote species collapse via hybridisation (Allendorf *et al*. 2001). Conservation approaches need to evaluate the role of hybridisation between populations and species as increased genetic variation may be supported by introgression (Bohling 2016).

Subspecies classifications have a major impact on the allocation of conservation resources (Zink 2004). Both *A. m. raglessi* and *A. m. indulkanna* are currently classified as subspecies based on plumage and morphological differences, and a mitochondrial divergence of 1.7% at ND2 (Black 2011; Austin *et al*. 2013). However, these subspecies are also known to have a continuous distribution and mitochondrial paraphyly (Slender *et al*. 2017). The lack of genetic differentiation and high level of gene flow between *A. m. raglessi* and *A. m. indulkanna* suggests these subspecies could be lumped into one Evolutionarily Significant Unit (ESU) for conservation purposes (Moritz 1994; Zink 2004). However, clinal genetic variation caused by IBD indicates subspecies classifications that require separate management approaches tailored specifically to *A. m. raglessi* or *A. m. indulkanna*. Other studies show that phenotypic variation moderately correlates with genotypic variation in natural populations (Wood *et al*. 2021). This supports the argument that morphologically divergent populations make a significant contribution to biodiversity. Managing the genetic variation captured by each of these subspecies will enable greater adaptive potential in the future (Fraser and Bernatchez 2001; Coates *et al*. 2018). Gene flow and genetic variation have an integral role in conservation management of subspecies, and sometimes units defined by morphotype is appropriate.

Speciation was traditionally thought to be more commonly associated with population divergence in allopatry, which has affected how we define species and subspecies (particularly those not in allopatry) (De Queiroz 2007; Marie Curie Speciation Network 2012). Examples where populations have undergone divergence with gene flow are now becoming more common since genomic techniques to assess gene flow are more accessible (Sousa and Hey 2013; Seehausen *et al*. 2014; Toews *et al*. 2016). This study shows that *A. m. raglessi* and *A. m. indulkanna* display patterns of morphological divergence that are in congruence with outlier loci associated with the subspecies occurrence in different vegetation types. This pattern is similar to other models of divergence with gene flow such as the little greenbul (*Andropadus virens*) where morphological divergence is more likely explained by the birds occurrence in different habitat types (savanna versus forest or mountain versus forest) than their allopatic history (Smith *et al*. 2005). In another model of divergence with gene flow (*Littorina saxatilis*), outlier loci have been genomically linked with loci that control phenotype, which are selected for according to ecotype (Hollander *et al*. 2015).

Lastly, Haenel *et al*. (2021) show how populations of the threespine stickleback fish (*Gasterosteus aculeatus*) that possess phenotypes associated with either stream or lake environments have developed reproductive isolation without any form of geographic barrier. This research outlines a mechanism for divergence with gene flow and suggests that the two parapatric TBGW subspecies in this study could be a model of divergence with gene flow that may continue to diverge in the future.

This study detected two interesting genomic patterns, the cause of which remain unresolved. Mitochondrial paraphyly at ND2 detected by Slender *et al*. (2017) predicted a low rate of genomic introgression as only 10% of *A. m. raglessi* individuals had an *A. m. indulkanna* haplotype. The contradictory results of this study based on nuclear markers could be explained by a number of processes, for example, selection for particular mtDNA haplotypes (Toews and Brelsford 2012; Morales *et al*. 2015; Morales *et al*. 2018), or greater dispersal of males compared to females. Other malurid species are known to display female sex-biased dispersal (Cockburn *et al*. 2003). However, the adult sample size per sex in this study was too small to investigate sex-biased dispersal. Intriguingly, this study also detected a third cluster of individuals in the middle of the *A. m. raglessi* range that displayed unique genomic variation. We can only hypothesize why these individuals were identified as distinct, but one possible scenario may be that limited gene flow between *A. m. raglessi* and another TBGW subspecies, such as *A. m. curnamona,* could be occurring or has more likely occurred in the past. The location of the nearest *A. m. curnamona* sighting is less than 100 km southeast from an *A. m. raglessi* sighting (Black *et al*. 2010). Further sampling of adult grasswrens and the inclusion of samples from other grasswren subspecies may reveal patterns of sex-biased dispersal and explain the source of distinct genomic variation detected within the *A. m. raglessi* population.

TBGW subspecies show asymmetric gene flow from *A. m. indulkanna* to *A. m. raglessi*. The dune field that runs between Lake Eyre and Lake Torrens demarcates the boundary of the asymmetry (Slender *et al*. 2017). Asymmetric gene flow could occur as the result of several processes, such as greater niche breadth in *A. m. indulkanna*, demographic or ecological differences on either side of the dune field that promote greater geneflow from *A. m. indulkanna* to *A. m. raglessi* (e.g. Oswald *et al*. 2017), or a mating advantage for *A. m. indulkanna* (e.g. Baldassarre and Webster 2013; Baldassarre *et al*. 2014; Slender *et al*. 2018b). The more frequent and intense response of *A. m. indulkanna* towards hetero-subspecific song compared to con-subspecific song could suggest *A. m. indulkanna* is more competitive than *A. m. raglessi*. *A. m. raglessi* did not show the same strength of response to hetero-subspecific song compared to con-subspecific song (Slender *et al*. 2018b). Further work is needed to test hypotheses regarding subspecies behaviour and habitat preference, landscape ecological productivity and stability, and mito-nuclear incompatibilities.

This study shows that two parapatric TBGW subspecies introgressed and that gene flow is asymmetric towards *A. m. raglessi*. Gene flow between the subspecies is limited by distance probably due to the low dispersing ability of the species as well as landscape heterogeneity. We suggest that these subspecies should be taxonomically (and administratively) managed as distinct units despite considerable introgression. Plant community type does not appear to limit geneflow nor does it provide a mechanism for increased genetic diversity at the parapatric margin as was predicted. However, adaptation to different plant community types suggests divergence with gene flow could be a pathway towards increased genomic variation in the future. This study provides an Australian arid zone example to show that gene flow between subspecies can increase genetic variation within a species. Increased gene flow is expected to facilitate persistence of the species through enhanced adaptive capacity. Populations that contain distinct genomic variation should be managed separately particularly in environments that are likely to be affected by future climate change.

## Data availability statement

The data that support this study will be shared upon reasonable request to the corresponding author.

## Acknowledgements

This research was conducted under the approval from the Department for Environment and Water who has also supplied the mapping vectors used in Figure 1. We thank the pastoral landowners and Arrium mining for access to their properties. We thank Dr. Andrew Black for advice on research outcomes and for insightful discussions about grasswren evolution and ecology. We thank Dr. Valeria Zanollo and all the volunteers that helped with data collection.

## References

Aguillon, S. M., Fitzpatrick, J. W., Bowman, R., Schoech, S. J., Clark, A. G., Coop, G., and Chen, N. (2017). Deconstructing isolation-by-distance: The genomic consequences of limited dispersal. PLoS Genetics 13(8), e1006911.

Aiello, B. R., Tan, M., Bin Sikandar, U., Alvey, A. J., Bhinderwala, B., Kimball, K. C., Barber, J. R., Hamilton, C. A., Kawahara, A. Y., and Sponberg, S. (2021). Adaptive shifts underlie the divergence in wing morphology in bombycoid moths. Proceedings of the Royal Society B: Biological Sciences 288(1956), 20210677.

Allan, J. R., Watson, J. E. M., Di Marco, M., O’Bryan, C. J., Possingham, H. P., Atkinson, S. C., and Venter, O. (2019). Hotspots of human impact on threatened terrestrial vertebrates. PLoS Biol 17(3), e3000158.

Allendorf, F. W., Leary, R. F., Spruell, P., and Wenburg, J. K. (2001). The problems with hybrids: setting conservation guidelines. Trends in Ecology and Evolution 16(11), 613–622.

Amos, J. N., Bennett, A. F., Mac Nally, R., Newell, G., Pavlova, A., Radford, J. Q., Thomson, J. R., White, M., and Sunnucks, P. (2012). Predicting landscape-genetic consequences of habitat loss, fragmentation and mobility for multiple species of woodland birds. Plos One 7(2), e30888.

Austin, J. J., Joseph, L., Pedler, L. P., and Black, A. B. (2013). Uncovering cryptic evolutionary diversity in extant and extinct populations of the southern Australian arid zone Western and Thick-billed Grasswrens (Passeriformes: Maluridae: *Amytornis*). Conservation Genetics 14(6), 1173–1184.

Baldassarre, D. T., and Webster, M. S. (2013). Experimental evidence that extra-pair mating drives asymmetrical introgression of a sexual trait. Proceedings of the Royal Society of London. Series B: Biological Sciences 280(1771), 20132175.

Baldassarre, D. T., White, T. A., Karubian, J., and Webster, M. S. (2014). Genomic and morphological analysis of a semipermeable avian hybrid zone suggests asymmetrical introgression of a sexual signal. Evolution 68(9), 2644–2657.

Balkenhol, N., Waits, L. P., and Dezzani, R. J. (2009). Statistical approaches in landscape genetics: an evaluation of methods for linking landscape and genetic data. Ecography 32(5), 818–830.

Berner, D., and Thibert-Plante, X. (2015). How mechanisms of habitat preference evolve and promote divergence with gene flow. J Evol Biol 28(9), 1641–1655.

Black, A. (2011). Subspecies of the Thick-billed Grasswren *Amytornis modestus* (Aves-Maluridae). Transactions of the Royal Society of South Australia 135(1), 26–38.

Black, A. (2016). Reappraisal of plumage and morphometic diversity in Thick-billed Grasswren *Amytornis modestus* (North, 1902), with description of a new subspecies. Bulletin of the British Ornithologists Club 136(1), 58-68.

Black, A., and Gower, P. (2017). ’Grasswrens: Australian outback identities.’ (Axiom: Stepney, South Australia.)

Black, A., Joseph, L., Pedler, L., and Carpenter, G. A. (2010). A taxonomic framework for interpreting evolution within the *Amytornis textilis-modestus* complex of grasswrens. Emu - Austral Ornithology 110(4), 358–363.

Black, A. B., Carpenter, G. A., and Pedler, L. P. (2011). Distribution and habitats of the Thick-Billed Grasswren *Amytornis modestus* and comparison with the Western Grasswren *Amytornis textilis myall* in South Australia. South Australian Ornithologist 37(2), 60–80.

Bohling, J. H. (2016). Strategies to address the conservation threats posed by hybridization and genetic introgression. Biological Conservation 203, 321–327.

Bradshaw, C. J. A. (2012). Little left to lose: deforestation and forest degradation in Australia since European colonization. Journal of Plant Ecology 5(1), 109-120.

Brandle, R. (1998) A biological survey of the Stony Deserts, South Australia, 1994-1997. Heritage and Biodiversity Section, Department for Environment, Heritage and Aboriginal Affairs, South Australia.

Case, T. J., and Taper, M. L. (2000). Interspecific competition, environmental gradients, gene flow, and the coevolution of species’ borders. The American Naturalist 155(5), 583–605.

Catchen, J., Hohenlohe, P. A., Bassham, S., Amores, A., and Cresko, W. A. (2013). Stacks: an analysis tool set for population genomics. Molecular Ecology 22(11), 3124–3140.

Chan, W. Y., Hoffmann, A. A., and Oppen, M. J. H. (2019). Hybridization as a conservation management tool. Conservation Letters 12(5).

Christidis, L., Rheindt, F. E., Boles, W. E., and Norman, J. A. (2010). Plumage patterns are good indicators of taxonomic diversity, but not of phylogenetic affinities, in Australian grasswrens Amytornis (Aves: Maluridae). Molecular Phylogenetics and Evolution 57(2), 868-877.

Christidis, L., Rheindt, F. E., Boles, W. E., and Norman, J. A. (2013). A re-appraisal of species diversity within the Australian grasswrens *Amytornis* (Aves: Maluridae). Australian Zoologist 36(4), 429–437.

Cicero, C. (2004). Barriers to sympatry between avian sibling species (Paridae: *Baeolophus*) in local secondary contact. Evolution 58(7), 1573–1587.

Coates, D. J., Byrne, M., and Moritz, C. (2018). Genetic Diversity and Conservation Units: Dealing With the Species-Population Continuum in the Age of Genomics. Frontiers in Ecology and Evolution 6.

Cockburn, A., Osmond, H. L., Mulder, R. A., Green, D. J., and Double, M. C. (2003). Divorce, dispersal and incest avoidance in the cooperatively breeding superb fairy-wren *Malurus cyaneus*. Journal of animal ecology 72, 189–202.

Davis, J. M. P., van Heerwaarden, B., Sgrò, C. M., Donald, J. A., and Kemp, D. J. (2013). Low genetic variation in cold tolerance linked to species distributions in butterflies. Evolutionary Ecology 28(3), 495–504.

De Queiroz, K. (2007). Species concepts and species delimitation. Systematic Biology 56(6), 879–886.

Deiner, K., Bik, H. M., Machler, E., Seymour, M., Lacoursiere-Roussel, A., Altermatt, F., Creer, S., Bista, I., Lodge, D. M., de Vere, N., Pfrender, M. E., and Bernatchez, L. (2017). Environmental DNA metabarcoding: Transforming how we survey animal and plant communities. Mol Ecol 26(21), 5872–5895.

Diniz-Filho, J. A. F., and Pires de Campos Telles, M. (2002). Spatial autocorrelation analysis and the identification of operational units for conservation in continuous populations. Conservation Biology 16(4), 924–935.

Dudaniec, R. Y., Yong, C. J., Lancaster, L. T., Svensson, E. I., and Hansson, B. (2018). Signatures of local adaptation along environmental gradients in a range-expanding damselfly (Ischnura elegans). Molecular Ecology 10.1111/mec.14709.

Dudgeon, C. L., Blower, D. C., Broderick, D., Giles, J. L., Holmes, B. J., Kashiwagi, T., Kruck, N. C., Morgan, J. A., Tillett, B. J., and Ovenden, J. R. (2012). A review of the application of molecular genetics for fisheries management and conservation of sharks and rays. J Fish Biol 80(5), 1789–1843.

Dupoué, A., Trochet, A., Richard, M., Sorlin, M., Guillon, M., Teulieres□Quillet, J., Vallé, C., Rault, C., Berroneau, M., Berroneau, M., Lourdais, O., Blaimont, P., Bertrand, R., Pottier, G., Calvez, O., Guillaume, O., Le Chevalier, H., Souchet, J., Le Galliard, J. F., Clobert, J., Aubret, F., and Razgour, O. (2020). Genetic and demographic trends from rear to leading edge are explained by climate and forest cover in a cold□adapted ectotherm. Diversity and Distributions 27(2), 267–281.

Earl, D. A., and vonHoldt, B. M. (2011). STRUCTURE HARVESTER: a website and program for visualizing STRUCTURE output and implementing the Evanno method. Conservation Genetics Resources 4(2), 359–361.

Elshire, R. J., Glaubitz, J. C., Sun, Q., Poland, J. A., Kawamoto, K., Buckler, E. S., and Mitchell, S. E. (2011). A robust, simple genotyping-by-sequencing (GBS) approach for high diversity species. PLoS One 6(5), e19379.

Evanno, G., Regnaut, S., and Goudet, J. (2005). Detecting the number of clusters of individuals using the software STRUCTURE: a simulation study. Molecular Ecology 14(8), 2611–2620.

Facelli, J. M., and Springbett, H. (2009). Why do some species in arid lands increase under grazing? Mechanisms that favour increased abundance of *Maireana pyramidata* in overgrazed chenopod shrublands of South Australia. Austral Ecology 34(5), 588–597.

Falush, D., Stephens, M., and Pritchard, J. K. (2003). Inference of population structure using multilocus genotype data: linked loci and correlated allele frequencies. Genetics 164, 1567–1587.

Fedorka, K. M., Winterhalter, W. E., Shaw, K. L., Brogan, W. R., and Mousseau, T. A. (2012). The role of gene flow asymmetry along an environmental gradient in constraining local adaptation and range expansion. Journal of Evolutionary Biology 25(8), 1676–1685.

Fischer, M. C., Rellstab, C., Leuzinger, M., Roumet, M., Gugerli, F., Shimizu, K. K., Holderegger, R., and Widmer, A. (2017). Estimating genomic diversity and population differentiation - an empirical comparison of microsatellite and SNP variation in Arabidopsis halleri. BMC Genomics 18(1), 69.

Flockhart, D. T., Pichancourt, J. B., Norris, D. R., and Martin, T. G. (2015). Unravelling the annual cycle in a migratory animal: breeding-season habitat loss drives population declines of monarch butterflies. J Anim Ecol 84(1), 155–165.

Foll, M., and Gaggiotti, O. (2008). A genome-scan method to identify selected loci appropriate for both dominant and codominant markers: a Bayesian perspective. Genetics 180(2), 977–993.

Forester, B. R., Jones, M. R., Joost, S., Landguth, E. L., and Lasky, J. R. (2016). Detecting spatial genetic signatures of local adaptation in heterogeneous landscapes. Mol Ecol 25(1), 104–120.

Forseth, T., Barlaup, B. T., Finstad, B., Fiske, P., Gjøsæter, H., Falkegård, M., Hindar, A., Mo, T. A., Rikardsen, A. H., Thorstad, E. B., Vøllestad, L. A., Wennevik, V., and Gibbs, M. (2017). The major threats to Atlantic salmon in Norway. ICES Journal of Marine Science 74(6), 1496–1513.

Frankham, R. (2010). Challenges and opportunities of genetic approaches to biological conservation. Biological Conservation 143(9), 1919–1927.

Fraser, D. J., and Bernatchez, L. (2001). Adaptive evolutionary conservation: towards a unified consept for defining conservation units. Molecular Ecology 10, 2741–2752.

Frichot, E., Schoville, S. D., Bouchard, G., and François, O. (2013). Testing for associations between loci and environmental gradients using latent factor mixed models. Mol Biol Evol 30(7), 1687-1699.

Gill, F., and Donsker, D. (2017) ’IOC World Bird List.’ Version 7.3. In Available at http://www.worldbirdnames.org/ [Verified 19 Sep 2017]

Gonçalves da Silva, A., Barendse, W., Kijas, J. W., Barris, W. C., McWilliam, S., Bunch, R. J., McCullough, R., Harrison, B., Hoelzel, A. R., and England, P. R. (2015). SNP discovery in nonmodel organisms: strand bias and base-substitution errors reduce conversion rates. Molecular Ecology Resources 15(4), 723–736.

Grant, P. R., and Grant, B. R. (2019). Hybridization increases population variation during adaptive radiation. Proc Natl Acad Sci U S A 116(46), 23216–23224.

Grismer, L. L. (2021). Comparative ecomorphology of the sandstone night lizard (Xantusia gracilis) and the granite night lizard (Xantusia henshawi). Vertebrate Zoology 71, 425–437.

Guillot, G., and Rousset, F. (2013). Dismantling the Mantel tests. Methods in Ecology and Evolution 4(4), 336–344.

Haenel, Q., Oke, K. B., Laurentino, T. G., Hendry, A. P., and Berner, D. (2021). Clinal genomic analysis reveals strong reproductive isolation across a steep habitat transition in stickleback fish. Nat Commun 12(1), 4850.

Hardy, O. J., and Vekemans, X. (2002). SPAGeDi: a versatile computer program to analyse spatial genetic structure at the individual or population levels. Molecular Ecoloty Notes 2, 618–620.

Harrisson, K. A., Pavlova, A., Amos, J. N., Takeuchi, N., Lill, A., Radford, J. Q., and Sunnucks, P. (2012). Fine-scale effects of habitat loss and fragmentation despite large-scale gene flow for some regionally declining woodland bird species. Landscape Ecology 27(6), 813–827.

Hoffman, A. A., and Blows, M. W. (1994). Species borders: ecological and evolutionary perspectives. Trends in Ecology and Evolution 9(6), 223–227.

Hohenlohe, P. A., Bassham, S., Etter, P. D., Stiffler, N., Johnson, E. A., and Cresko, W. A. (2010). Population genomics of parallel adaptation in threespine stickleback using sequenced RAD tags. PLoS Genetics 6(2), e1000862.

Hollander, J., Galindo, J., and Butlin, R. K. (2015). Selection on outlier loci and their association with adaptive phenotypes in *Littorina saxatilis* contact zones. Journal of Evolutionary Biology 28(2), 328–337.

Jakobsson, M., and Rosenberg, N. A. (2007). CLUMPP: a cluster matching and permutation program for dealing with label switching and multimodality in analysis of population structure. Bioinformatics 23(14), 1801–1806.

Jellinek, S., Harrison, P. A., Tuck, J., and Te, T. (2020). Replanting agricultural landscapes: how well do plants survive after habitat restoration? Restoration Ecology 28(6), 1454–1463.

Jenkins, N., and Hoffman, A. A. (1999). Limits to the Southern Border of Drosophila serrata: Cold Resistance, Heritable Variation, and Trade-Offs. Evolution 53(6), 1823–1834.

Jessop, P. (1995) Response of arid vegetation to cattle grazing and the development of indicators for range assessment with particular reference to the rangelands of northern South Australia. MASc Thesis, Adelaide University, Adelaide

Jombart, T. (2008). adegenet: a R package for the multivariate analysis of genetic markers. Bioinformatics 24(11), 1403–1405.

Jombart, T., Devillard, S., and Balloux, F. (2010). Discriminant analysis of principal components: a new method for the analysis of genetically structured populations. BMC Genetics 11, 94.

Kierepka, E. M., and Latch, E. K. (2015). Performance of partial statistics in individual-based landscape genetics. Mol Ecol Resour 15(3), 512–525.

Kingsford, R. T. (2000). Ecological impacts of dams, water diversions and river management on floodplains wetlands in Australia. Austral Ecology 25, 109–127.

Kirkpatrick, M., and Barton, N. H. (1997). Evolution of species’ range. The American Naturalist 150(1), 1–23.

Koizumi, I., Yamamoto, S., and Maekawa, K. (2006). Decomposed pairwise regression analysis of genetic and geographic distances reveals a metapopulation structure of stream-dwelling Dolly Varden charr. Mol Ecol 15(11), 3175–3189.

Lenormand, T. (2002). Gene flow and the limits to natural selection. Trends in Ecology and Evolution 17(4), 183–189.

Lindenmayer, D. B., and Burgman, M. (2005). ’Practical Conservation Biology.’ (CSIRO Publishing: Melbourne.)

Lioubimtseva, E. (2004). Climate change in arid environments: revisiting the past to understand the future. Progress in Physical Geography 28(4), 502–530.

Lischer, H. E., and Excoffier, L. (2012). PGDSpider: an automated data conversion tool for connecting population genetics and genomics programs. Bioinformatics 28(2), 298–299.

Lu, F., Lipka, A. E., Glaubitz, J., Elshire, R., Cherney, J. H., Casler, M. D., Buckler, E. S., and Costich, D. E. (2013). Switchgrass genomic diversity, ploidy, and evolution: novel insights from a network-based SNP discovery protocol. PLoS Genetics 9(1), e1003215.

Mallen-Cooper, M., and Zampatti, B. P. (2020). Restoring the ecological integrity of a dryland river: Why low flows in the Barwon–Darling River must flow. Ecological Management & Restoration 21(3), 218–228.

Meirmans, P. G. (2012). The trouble with isolation by distance. Molecular Ecology 21, 2839–2846.

Meirmans, P. G. (2020). genodive version 3.0: Easy-to-use software for the analysis of genetic data of diploids and polyploids. Molecular Ecology Resources 20(4), 1126–1131.

Meirmans, P. G., and Van Tienderen, P. H. (2004). genotype and genodive: two programs for the analysis of genetic diversity of asexual organisms. Molecular Ecology Notes 4(4), 792–794.

Moerman, F., Fronhofer, E. A., Wagner, A., and Altermatt, F. (2020). Gene swamping alters evolution during range expansions in the protist Tetrahymena thermophila. Biol Lett 16(6), 20200244.

Morales, H. E., Pavlova, A., Amos, N., Major, R., Kilian, A., Greening, C., and Sunnucks, P. (2018). Concordant divergence of mitogenomes and a mitonuclear gene cluster in bird lineages inhabiting different climates. Nat Ecol Evol2018/07/11 10.1038/s41559-018-0606-3.

Morales, H. E., Pavlova, A., Joseph, L., and Sunnucks, P. (2015). Positive and purifying selection in mitochondrial genomes of a bird with mitonuclear discordance. Molecular Ecology 24(11), 2820–2837.

Moritz, C. (1994). Defining Evolutionary Significant Units for conservation. Trends in Ecology and Evolution 9, 373–375.

Mynhardt, S., Bennett, N. C., and Bloomer, P. (2020). New insights from RADseq data on differentiation in the Hottentot golden mole species complex from South Africa. Mol Phylogenet Evol 143, 106667.

Navarro, T., Alados, C. L., and Cabezudo, B. (2006). Changes in plant functional types in response to goat and sheep grazing in two semi-arid shrublands of SE Spain. Journal of Arid Environments 64(2), 298–322.

Nei, M. (1977). F-statistics and analysis of gene diversity in subdivided populations. Annals of Human Genetics 41(2), 225–233.

Network, M. C. S. (2012). What do we need to know about speciation? Trends in Ecology and Evolution 27(1), 27–39.

Newbold, T., Hudson, L. N., Hill, S. L., Contu, S., Lysenko, I., Senior, R. A., Borger, L., Bennett, D. J., Choimes, A., Collen, B., Day, J., De Palma, A., Diaz, S., Echeverria-Londono, S., Edgar, M. J., Feldman, A., Garon, M., Harrison, M. L., Alhusseini, T., Ingram, D. J., Itescu, Y., Kattge, J., Kemp, V., Kirkpatrick, L., Kleyer, M., Correia, D. L., Martin, C. D., Meiri, S., Novosolov, M., Pan, Y., Phillips, H. R., Purves, D. W., Robinson, A., Simpson, J., Tuck, S. L., Weiher, E., White, H. J., Ewers, R. M., Mace, G. M., Scharlemann, J. P., and Purvis, A. (2015). Global effects of land use on local terrestrial biodiversity. Nature 520(7545), 45–50.

Norman, J. A., and Christidis, L. (2016). Ecological opportunity and the evolution of habitat preferences in an arid-zone bird: implications for speciation in a climate-modified landscape. Scientific Reports 6, 19613.

Nowicki, S., and Searcy, W. A. (2005). Song and Mate Choice in Birds: How the Development of Behavior Helps Us Understand Function. The Auk 122(1), 1.

Oswald, J. A., Overcast, I., Mauck III, W. M., Andersen, M. J., and Smith, B. T. (2017). Isolation with asymmetric gene flow during the nonsynchronous divergence of dry forest birds. Molecular Ecology 26, 1386–1400.

Paparella, S., Araujo, S. S., Rossi, G., Wijayasinghe, M., Carbonera, D., and Balestrazzi, A. (2015). Seed priming: state of the art and new perspectives. Plant Cell Rep 34(8), 1281–1293.

Paradis, E. (2010). pegas: an R package for population genetics with an integrated-modular approach. Bioinformatics 26(3), 419–420.

Peakall, R., and Smouse, P. E. (2012). GenAlEx 6.5: genetic analysis in Excel. Population genetic software for teaching and research--an update. Bioinformatics 28(19), 2537–2539.

Peakall, R. O. D., and Smouse, P. E. (2006). GenAlEx 6: genetic analysis in Excel. Population genetic software for teaching and research. Molecular Ecology Notes 6(1), 288–295.

Perez, M. F., Franco, F. F., Bombonato, J. R., Bonatelli, I. A. S., Khan, G., Romeiro-Brito, M., Fegies, A. C., Ribeiro, P. M., Silva, G. A. R., Moraes, E. M., and Burridge, C. (2018). Assessing population structure in the face of isolation by distance: Are we neglecting the problem? Diversity and Distributions 24(12), 1883–1889.

Pickup, G. (1998). Desertification and climate change - the Australian perspective. Climate Research 11, 51–63.

Poland, J. A., Brown, P. J., Sorrells, M. E., and Jannink, J.-L. (2012). Development of high-density genetic maps for Barley and Wheat using a novel two-enzyme genotyping-by-sequencing approach. PLoS ONE 7(2), e32253.

Pritchard, J. K., Stephens, M., and Donnelly, P. (2000). Inference of population structure using multilocus genotype data. Genetics 155(2), 945–959.

Purcell, S., Neale, B., Todd-Brown, K., Thomas, L., Ferreira, M. A., Bender, D., Maller, J., Sklar, P., de Bakker, P. I., Daly, M. J., and Sham, P. C. (2007). PLINK: a tool set for whole-genome association and population-based linkage analyses. The American Journal of Human Genetics 81(3), 559–575.

Rambaut, A., Drummond, A. J., Xie, D., Baele, G., and Suchard, M. A. (2015) Tracer v1.6. http://beast.community

Reynolds, R. G. (2011) Islands Metapopulations and Archipelagos: Genetic Equilibrium and Non-Equilibrium Dynamics of Structured Populations in teh Context of Conservation., University of Tennessee,

Rosauer, D. F., Byrne, M., Blom, M. P. K., Coates, D. J., Donnellan, S., Doughty, P., Keogh, J. S., Kinloch, J., Laver, R. J., Myers, C., Oliver, P. M., Potter, S., Rabosky, D. L., Afonso Silva, A. C., Smith, J., and Moritz, C. (2018). Real-world conservation planning for evolutionary diversity in the Kimberley, Australia, sidesteps uncertain taxonomy. Conservation Letters 10.1111/conl.12438.

Rosenberg, N. A. (2004). DISTRUCT: a program for the graphical display of population structure. Molecular Ecology Notes 4(1), 137–138.

Rossetto, M., Yap, J.-Y. S., Lemmon, J., Bain, D., Bragg, J., Hogbin, P., Gallagher, R., Rutherford, S., Summerell, B., and Wilson, T. C. (2021). A conservation genomics workflow to guide practical management actions. Global Ecology and Conservation 26, e01492.

Rousset, F. (1997). Genetic differentiation and estimation of gene flow from *F*-Statistics under isolation by distance. Genetics 145, 1219–1228.

Rousset, F. (2000). Genetic differentiation between individuals. Journal of Evolutionary Biology 13, 58–62.

Saccheri, I., Kuussaari, M., Kankare, M., Vikman, P., Fortelius, W., and Hanski, I. (1998). Inbreeding and extinction in a butterfly metapopulation. Nature 392, 491–494.

Sanchez-Guillen, R. A., Cordoba-Aguilar, A., Hansson, B., Ott, J., and Wellenreuther, M. (2016). Evolutionary consequences of climate-induced range shifts in insects. Biol Rev Camb Philos Soc 91(4), 1050–1064.

Seehausen, O., Butlin, R. K., Keller, I., Wagner, C. E., Boughman, J. W., Hohenlohe, P. A., Peichel, C. L., Saetre, G.-P., Bank, C., Brännström, A., Brelsford, A., Clarkson, C. S., Eroukhmanoff, F., Feder, J. L., Fischer, M. C., Foote, A. D., Franchini, P., Jiggins, C. D., Jones, F. C., Lindholm, A. K., Lucek, K., Maan, M. E., Marques, D. A., Martin, S. H., Matthews, B., Meier, J. I., Möst, M., Nachman, M. W., Nonaka, E., Rennison, D. J., Schwarzer, J., Watson, E. T., Westram, A. M., and Widmer, A. (2014). Genomics and the origin of species. Nature Reviews. Genetics 15(3), 176–192.

Skroblin, A., and Murphy, S. (2013). The conservation status of Australian malurids and their value as models in understanding land-management issues. Emu - Austral Ornithology 113(3), 309–318.

Slatyer, R. O. (1961). Methodology of a water balance study conducted on a desert woodland (*Acacia anuera* F.Muell) community in central Australia. UNESCO Arid Zone Research 16, 15–26.

Slender, A. L., Louter, M., Gardner, M. G., and Kleindorfer, S. (2017). Patterns of morphological and mitochondrial diversity in parapatric subspecies of the Thick-billed Grasswren (*Amytornis modestus*). Emu - Austral Ornithology 117(3), 264–275.

Slender, A. L., Louter, M., Gardner, M. G., and Kleindorfer, S. (2018a). Plant community predicts the distribution and occurrence of thick-billed grasswren subspecies (*Amytornis modestus*) in a region of parapatry. Australian Journal of Zoology 65(4), 273-282.

Slender, A. L., Louter, M., Gardner, M. G., and Kleindorfer, S. (2018b). Thick-billed grasswren (*Amytornis modestus*) songs differ across subspecies and elicit different subspecific behavioural responses. Transactions of the Royal Society of South Australia 142(2), 105–121.

Smith, L. M., and Burgoyne, L. A. (2004). Collecting, archiving and processing DNA from wildlife samples using FTA® databasing paper. BMC Ecology 4, 4.

Smith, T. B., Calsbeek, R., Wayne, R. K., Holder, K. H., Pires, D., and Bardeleben, C. (2005). Testing alternative mechanisms of evolutionary divergence in an African rain forest passerine bird. Journal of Evolutionary Biology 18(2), 257–268.

Smouse, P. E., and Peakall, R. (1999). Spatial autocorrelation analysis of individual multiallele and multilocus genetic structure. Heredity 82, 561–573.

Sousa, V., and Hey, J. (2013). Understanding the origin of species with genome-scale data: modelling gene flow. Nat Rev Genet 14(6), 404–414.

Steiner, C. C., Putnam, A. S., Hoeck, P. E. A., and Ryder, O. A. (2013). Conservation genomics of threatened animal species. Annual Review of Animal Biosciences 1(1), 261–281.

Thompson, T. Q., Bellinger, M. R., O’Rourke, S. M., Prince, D. J., Stevenson, A. E., Rodrigues, A. T., Sloat, M. R., Speller, C. F., Yang, D. Y., Butler, V. L., Banks, M. A., and Miller, M. R. (2019). Anthropogenic habitat alteration leads to rapid loss of adaptive variation and restoration potential in wild salmon populations. Proc Natl Acad Sci U S A 116(1), 177–186.

Toews, D. P. L., and Brelsford, A. (2012). The biogeography of mitochondrial and nuclear discordance in animals. Molecular Ecology 21(16), 3907–3930.

Toews, D. P. L., Campagna, L., Taylor, S. A., Balakrishnan, C. N., Baldassarre, D. T., Deane-Coe, P. E., Harvey, M. G., Hooper, D. M., Irwin, D. E., Judy, C. D., Mason, N. A., McCormack, J. E., McCracken, K. G., Oliveros, C. H., Safran, R. J., Scordato, E. S. C., Stryjewski, K. F., Tigano, A., Uy, J. A. C., and Winger, B. M. (2016). Genomic approaches to understanding population divergence and speciation in birds. The Auk 133(1), 13–30.

Vaghefi, S. A., Keykhai, M., Jahanbakhshi, F., Sheikholeslami, J., Ahmadi, A., Yang, H., and Abbaspour, K. C. (2019). The future of extreme climate in Iran. Sci Rep 9(1), 1464.

van Strien, M. J., Holderegger, R., and Van Heck, H. J. (2015). Isolation-by-distance in landscapes: considerations for landscape genetics. Heredity 114(1), 27–37.

Wang, J. (2011). COANCESTRY: a program for simulating, estimating and analysing relatedness and inbreeding coefficients. Molecular Ecology Resources 11(1), 141–145.

Warren, W. C., Clayton, D. F., Ellegren, H., Arnold, A. P., Hillier, L. W., Kunstner, A., Searle, S., White, S., Vilella, A. J., Fairley, S., Heger, A., Kong, L., Ponting, C. P., Jarvis, E. D., Mello, C. V., Minx, P., Lovell, P., Velho, T. A., Ferris, M., Balakrishnan, C. N., Sinha, S., Blatti, C., London, S. E., Li, Y., Lin, Y. C., George, J., Sweedler, J., Southey, B., Gunaratne, P., Watson, M., Nam, K., Backstrom, N., Smeds, L., Nabholz, B., Itoh, Y., Whitney, O., Pfenning, A. R., Howard, J., Volker, M., Skinner, B. M., Griffin, D. K., Ye, L., McLaren, W. M., Flicek, P., Quesada, V., Velasco, G., Lopez-Otin, C., Puente, X. S., Olender, T., Lancet, D., Smit, A. F., Hubley, R., Konkel, M. K., Walker, J. A., Batzer, M. A., Gu, W., Pollock, D. D., Chen, L., Cheng, Z., Eichler, E. E., Stapley, J., Slate, J., Ekblom, R., Birkhead, T., Burke, T., Burt, D., Scharff, C., Adam, I., Richard, H., Sultan, M., Soldatov, A., Lehrach, H., Edwards, S. V., Yang, S. P., Li, X., Graves, T., Fulton, L., Nelson, J., Chinwalla, A., Hou, S., Mardis, E. R., and Wilson, R. K. (2010). The genome of a songbird. Nature 464(7289), 757–762.

Wellborn, G. A., and Langerhans, R. B. (2015). Ecological opportunity and the adaptive diversification of lineages. Ecology and Evolution 5(1), 176–195.

Whiteley, A. R., Fitzpatrick, S. W., Funk, W. C., and Tallmon, D. A. (2015). Genetic rescue to the rescue. Trends Ecol Evol 30(1), 42–49.

Whitlock, M. C., and Lotterhos, K. E. (2015). Reliable Detection of Loci Responsible for Local Adaptation: Inference of a Null Model through Trimming the Distribution of F(ST). Am Nat 186 Suppl 1, S24–36.

Williams, O. B. (1982) The vegetation of arid Australia: a biotic appraisal. In ’Evolution of the flora and fauna of arid Australia.’ (Eds. WR Barker and PJM Greenslade) pp. 3–14. (Peacock Press: Adelaide)

Willoughby, J. R., Sundaram, M., Wijayawardena, B. K., Kimble, S. J. A., Ji, Y., Fernandez, N. B., Antonides, J. D., Lamb, M. C., Marra, N. J., and DeWoody, J. A. (2015). The reduction of genetic diversity in threatened vertebrates and new recommendations regarding IUCN conservation rankings. Biological Conservation 191, 495–503.

Wilson, G. A., and Rannala, B. (2003). Bayesian inference of recent migration rates using multilocus genotypes. Genetics 163, 1177–1191.

Woinarski, J. C., Burbidge, A. A., and Harrison, P. L. (2015). Ongoing unraveling of a continental fauna: decline and extinction of Australian mammals since European settlement. Proceedings of the National Academy of Sciences 112(15), 4531–4540.

Wood, Z. T., Wiegardt, A. K., Barton, K. L., Clark, J. D., Homola, J. J., Olsen, B. J., King, B. L., Kovach, A. I., and Kinnison, M. T. (2021). Meta analysis: Congruence of genomic and phenotypic differentiation across diverse natural study systems. Evolutionary Applications 10.1111/eva.13264.

Zink, R. M. (2004). The role of subspecies in obscuring avian biological diversity and misleading conservation policy. Proceedings of the Royal Society of London. Series B: Biological Sciences 271(1539), 561–564.

