## supporting information for "Gene flow between two thick-billed grasswren subspecies with low dispersal creates a genomic pattern of isolation-by-distance"

**Table S1.** List of samples used in the study. Samples listed in the same order as in Figure 3. ND = Not Done, MS = Museum Sample and CS = Contemporary Sample. FTA = Whatmann FTA card (Sigma-Aldrich Pty Ltd, NSW, Australia), Blood in solution (Seutin et al. 1991), tissue samples were stored in ethanol, DNA extracts were from Austin et al. (2013).

| Sample | Year | Sample Collection | Sample Type | Zone | Locality | Locality abbreviation | Sex | Haplogroup |
| --- | --- | --- | --- | --- | --- | --- | --- | --- |
| SAMA BS7001 | 2008 | MS | Tissue | A | Oodnadattta West | OOW | Unknown | Western |
| ABBBS 03666538 | 2013 | CS | Blood in solution | A | Oodnadattta West | OOW | Female | Western |
| ABBBS 03666537 | 2013 | CS | Blood in solution | A | Oodnadattta West | OOW | Male | Western |
| ABBBS 03666541 | 2013 | CS | Blood in solution | A | Oodnadattta West | OOW | Female | Western |
| ABBBS 03666540 | 2013 | CS | Blood in solution | A | Oodnadattta West | OOW | Female | Western |
| ABBBS 03666539 | 2013 | CS | Blood in solution | A | Oodnadattta West | OOW | Male | Western |
| ABBBS 03666534 | 2013 | CS | Blood in solution | A | Oodnadattta West | OOW | Male | Western |
| SAMA B55706 | 2007 | MS | DNA extract | A | Oodnadattta East | OOE | Female | Western |
| SAMA B55705 | 2007 | MS | DNA extract | A | Oodnadattta East | OOE | Female | Western |
| SAMA B55707 | 2007 | MS | DNA extract | A | Mount Barry station | MTB | Female | Western |
| SAMA B55669 | 2007 | MS | DNA extract | A | Coober Pedy | COP | Male | Western |
| ABBBS 03666570 | 2013 | CS | Blood in solution | A | Coober Pedy | COP | Male | Western |
| ABBBS 03666555 | 2013 | CS | Blood in solution | A | Coober Pedy | COP | Male | Western |
| ABBBS 03666554 | 2013 | CS | Blood in solution | A | Coober Pedy | COP | Female | Western |
| ABBBS 03666574 | 2013 | CS | Blood in solution | A | Coober Pedy | COP | Male | Western |
| ABBBS 03666573 | 2013 | CS | Blood in solution | A | Coober Pedy | COP | Female | Western |
| ABBBS 03666572 | 2013 | CS | Blood in solution | A | Coober Pedy | COP | Female | Western |
| ABBBS 03666571 | 2013 | CS | Blood in solution | A | Coober Pedy | COP | Male | Western |
| ABBBS 03666562 | 2013 | CS | Blood in solution | A | Coober Pedy | COP | Male | Western |
| ABBBS 03666557 | 2013 | CS | Blood in solution | A | Coober Pedy | COP | Male | Western |
| ABBBS 03666560 | 2013 | CS | Blood in solution | A | Peculiar Knob | PEK | Female | Western |
| ABBBS 03666559 | 2013 | CS | Blood in solution | A | Peculiar Knob | PEK | Male | Western |
| ABBBS 03666558 | 2013 | CS | Blood in solution | A | Peculiar Knob | PEK | Female | Western |
| ABBBS 03666566 | 2013 | CS | Blood in solution | A | Peculiar Knob | PEK | Female | Western |
| ABBBS 03666561 | 2013 | CS | Blood in solution | A | Peculiar Knob | PEK | Male | Western |
| ABBBS 03666568 | 2013 | CS | Blood in solution | A | Peculiar Knob | PEK | Female | Western |
| SAMA B55670 | 2007 | MS | DNA extract | A | Peculiar Knob | PEK | Male | Western |
| ABBBS 03666546 | 2013 | CS | Blood in solution | A | William Creek | WIC | Male | Western |
| ABBBS 03666543 | 2013 | CS | Blood in solution | A | William Creek | WIC | Male | Western |
| ABBBS 03666548 | 2013 | CS | Blood in solution | A | William Creek | WIC | Female | Western |
| ABBBS 03666552 | 2013 | CS | Blood in solution | A | William Creek | WIC | Female | Western |
| ABBBS 03666551 | 2013 | CS | Blood in solution | A | William Creek | WIC | Male | Western |
| ABBBS 03666547 | 2013 | CS | Blood in solution | A | William Creek | WIC | Male | Western |
| SAMA B59006* | 2013 | CS | Blood in solution | A | Coward Springs Railway Siding | COS | Female | Western |
| ABBBS 03666582 | 2013 | CS | Blood in solution | A | Coward Springs Railway Siding | COS | Male | Western |
| ABBBS 03666577 | 2013 | CS | Blood in solution | A | Coward Springs Railway Siding | COS | Male | Western |
| ABBBS 03666585 | 2013 | CS | Blood in solution | A | Coward Springs Railway Siding | COS | Male | Western |
| ABBBS 03666584 | 2013 | CS | Blood in solution | A | Coward Springs Railway Siding | COS | Female | Western |
| ABBBS 03666580 | 2013 | CS | FTA | A | Coward Springs Railway Siding | COS | Male | Western |
| ABBBS 03666579 | 2013 | CS | Blood in solution | A | Coward Springs Railway Siding | COS | Female | Western |
| ABBBS 03666576 | 2013 | CS | Blood in solution | A | Coward Springs Railway Siding | COS | Male | Western |
| ABBBS 03666575 | 2013 | CS | Blood in solution | A | Coward Springs Railway Siding | COS | Female | Western |
| SAMA B59004* | 2013 | CS | Blood in solution | A | Coward Springs Railway Siding | COS | Male | Western |
| SAMA B59005* | 2013 | CS | Blood in solution | A | Coward Springs Railway Siding | COS | Male | Western |
| SAMA B55668 | 2007 | MS | DNA extract | AB | Mulgaria station | MUL | Female | Western |
| SAMA B55667 | 2007 | MS | DNA extract | AB | Mulgaria station | MUL | Male | Western |
| SAMA B59003* | 2013 | CS | Blood in solution | AB | Mulgaria station | MUL | Male | Western |
| ABBBS 03666530 | 2014 | CS | Blood in solution | AB | Stuart Creek station | STC | Male | Western |
| ABBBS 03666527 | 2014 | CS | Blood in solution | AB | Stuart Creek station | STC | Female | Eastern |
| ABBBS 03666529 | 2014 | CS | Blood in solution | AB | Stuart Creek station | STC | Female | Eastern |
| ABBBS 03666528 | 2014 | CS | Blood in solution | AB | Stuart Creek station | STC | Female | Western |
| ABBBS 03666532 | 2014 | CS | Blood in solution | AB | Stuart Creek station | STC | Female | Eastern |
| ABBBS 03666518 | 2014 | CS | Blood in solution | AB | Stuart Creek station | STC | Male | Eastern |
| ABBBS 03666523 | 2014 | CS | Blood in solution | AB | Stuart Creek station | STC | Male | Western |
| ABBBS 03666522 | 2014 | CS | Blood in solution | AB | Stuart Creek station | STC | Female | Eastern |
| ABBBS 03666526 | 2014 | CS | Blood in solution | AB | Stuart Creek station | STC | Female | Eastern |
| ABBBS 03666524 | 2014 | CS | Blood in solution | AB | Stuart Creek station | STC | Male | Eastern |
| ABBBS 03667351 | 2013 | CS | Blood in solution | B | Witchelina Nature Reserve | WIT | Female | Eastern |
| ABBBS 03667352 | 2013 | CS | Blood in solution | B | Witchelina Nature Reserve | WIT | Male | Eastern |
| ABBBS 03667396 | 2013 | CS | Blood in solution | B | Witchelina Nature Reserve | WIT | Male | Eastern |
| ABBBS 03667397 | 2013 | CS | Blood in solution | B | Witchelina Nature Reserve | WIT | Female | Western |
| ABBBS 03666887 | 2013 | CS | Blood in solution | B | Witchelina Nature Reserve | WIT | Female | Eastern |
| ABBBS 03666886 | 2013 | CS | Blood in solution | B | Witchelina Nature Reserve | WIT | Male | Western |
| ABBBS 02572570 | 2012 | CS | FTA | B | Witchelina Nature Reserve | WIT | Male | Eastern |
| ABBBS 03666589 | 2013 | CS | Blood in solution | B | Witchelina Nature Reserve | WIT | Male | Western |
| ABBBS 03666593 | 2014 | CS | Blood in solution | B | Witchelina Nature Reserve | WIT | Male | Eastern |
| ABBBS 03667362 | 2013 | CS | Blood in solution | B | Witchelina Nature Reserve | WIT | Male | Eastern |
| ABBBS 03667393 | 2013 | CS | Blood in solution | B | Witchelina Nature Reserve | WIT | Male | Western |
| ABBBS 03667395 | 2013 | CS | Blood in solution | B | Witchelina Nature Reserve | WIT | Female | Eastern |
| ABBBS 03666899 | 2013 | CS | Blood in solution | B | Witchelina Nature Reserve | WIT | Male | Eastern |
| ABBBS 03667367 | 2013 | CS | Blood in solution | B | Witchelina Nature Reserve | WIT | Female | Eastern |
| ABBBS 03667368 | 2013 | CS | Blood in solution | B | Witchelina Nature Reserve | WIT | Male | Eastern |
| ABBBS 03666888 | 2013 | CS | Blood in solution | B | Witchelina Nature Reserve | WIT | Female | Eastern |
| NestT36 | 2013 | CS | Tissue | B | Witchelina Nature Reserve | WIT | Male | Western |
| ABBBS 03667387 | 2013 | CS | Blood in solution | B | Witchelina Nature Reserve | WIT | Female | Eastern |
| ABBBS 03666875 | 2013 | CS | FTA | B | Witchelina Nature Reserve | WIT | Female | Western |
| ABBBS 03666876 | 2013 | CS | FTA | B | Witchelina Nature Reserve | WIT | Male | Eastern |
| ABBBS 03636691 | 2014 | CS | Blood in solution | B | Witchelina Nature Reserve | WIT | Female | Western |
| ABBBS 03636688 | 2014 | CS | Blood in solution | B | Witchelina Nature Reserve | WIT | Male | Eastern |
| ABBBS 03636690 | 2014 | CS | Blood in solution | B | Witchelina Nature Reserve | WIT | Female | Eastern |
| ABBBS 03636697 | 2014 | CS | Blood in solution | B | Witchelina Nature Reserve | WIT | Male | Eastern |
| ABBBS 03667374 | 2013 | CS | Blood in solution | B | Witchelina Nature Reserve | WIT | Female | Eastern |
| ABBBS 03667371 | 2013 | CS | Blood in solution | B | Witchelina Nature Reserve | WIT | Male | Eastern |
| ABBBS 03667373 | 2013 | CS | Blood in solution | B | Witchelina Nature Reserve | WIT | Female | Eastern |
| ABBBS 03666590 | 2013 | CS | Blood in solution | B | Witchelina Nature Reserve | WIT | Male | Eastern |
| ABBBS 03667389 | 2013 | CS | Blood in solution | B | Witchelina Nature Reserve | WIT | Female | Eastern |
| ABBBS 03667383 | 2013 | CS | Blood in solution | B | Witchelina Nature Reserve | WIT | Female | Eastern |
| ABBBS 03667384 | 2013 | CS | Blood in solution | B | Witchelina Nature Reserve | WIT | Male | Western |
| ABBBS 03636696 | 2014 | CS | Blood in solution | B | Witchelina Nature Reserve | WIT | Male | Eastern |
| ABBBS 02572568 | 2013 | CS | Blood in solution | B | Witchelina Nature Reserve | WIT | Male | Eastern |
| ABBBS 03666891 | 2013 | CS | FTA | B | Witchelina Nature Reserve | WIT | Male | Eastern |
| ABBBS 03666893 | 2013 | CS | FTA | B | Witchelina Nature Reserve | WIT | Female | Eastern |
| ABBBS 02572572 | 2012 | CS | FTA | B | Witchelina Nature Reserve | WIT | Male | Eastern |
| ABBBS 03666897 | 2013 | CS | Blood in solution | B | Witchelina Nature Reserve | WIT | Female | Eastern |
| ABBBS 03666898 | 2013 | CS | Blood in solution | B | Witchelina Nature Reserve | WIT | Male | Eastern |
| ABBBS 03667363 | 2013 | CS | Blood in solution | B | Witchelina Nature Reserve | WIT | Male | Eastern |
| ABBBS 03666588 | 2013 | CS | Blood in solution | B | Witchelina Nature Reserve | WIT | Male | Eastern |
| ABBBS 03667347 | 2014 | CS | Blood in solution | B | Witchelina Nature Reserve | WIT | Female | Eastern |
| ABBBS 03666586 | 2013 | CS | Blood in solution | B | Witchelina Nature Reserve | WIT | Female | Eastern |
| SAMA B59025* | 2014 | CS | Tissue | B | Witchelina Nature Reserve | WIT | Male | Eastern |
| ABBBS 03667334 | 2014 | CS | Blood in solution | B | Witchelina Nature Reserve | WIT | Female | Eastern |
| ABBBS 03667332 | 2014 | CS | Blood in solution | B | Witchelina Nature Reserve | WIT | Male | Eastern |
| ABBBS 03667327 | 2014 | CS | Blood in solution | B | Witchelina Nature Reserve | WIT | Male | Eastern |
| ABBBS 03667331 | 2014 | CS | FTA | B | Witchelina Nature Reserve | WIT | Female | Eastern |
| ABBBS 03667381 | 2013 | CS | Blood in solution | B | Witchelina Nature Reserve | WIT | Male | Western |
| ABBBS 03636681 | 2014 | CS | Blood in solution | B | Witchelina Nature Reserve | WIT | Male | Eastern |
| ABBBS 03666882 | 2013 | CS | Blood in solution | B | Witchelina Nature Reserve | WIT | Male | Eastern |
| ABBBS 03667365 | 2013 | CS | Blood in solution | B | Witchelina Nature Reserve | WIT | Male | Eastern |
| ANWC B40189 | 1985 | MS | DNA extract | B | Witchelina Nature Reserve | WIT | Male | Eastern |
| ANWC B40190 | 1985 | MS | DNA extract | B | Witchelina Nature Reserve | WIT | Female | Eastern |
| ANWC B40177 | 1985 | MS | DNA extract | B | Mount Lyndhurst station | MTL | Female | Eastern |
| ANWC B40180 | 1985 | MS | DNA extract | B | Mount Lyndhurst station | MTL | Male | Eastern |
| ABBBS 03666512 | 2014 | CS | Blood in solution | B | Mount Lyndhurst station | MTL | Male | Eastern |
| ABBBS 03666511 | 2014 | CS | Blood in solution | B | Mount Lyndhurst station | MTL | Female | Eastern |
| ABBBS 03666514 | 2014 | CS | Blood in solution | B | Mount Lyndhurst station | MTL | Male | Eastern |
| ABBBS 03666513 | 2014 | CS | Blood in solution | B | Mount Lyndhurst station | MTL | Female | Eastern |
| SAMA B55666 | 2007 | MS | DNA extract | B | Murnpeowie station | MUR | Male | Eastern |
| SAMA B56154 | 2009 | MS | DNA extract | B | Murnpeowie station | MUR | Female | Eastern |

*Samples subsequently submitted to the South Australian museum after sample collection for this study.

**Table S2**. Results of discontiguous megablast for sequence similarity between outlier loci and the Zebra Finch (*Taeniopygia guttata*) genome. E – value is the expected number of hits at random, PI is the percent identity, PREDICTED means that the protein translations from the Zebra Finch sequences have not been tested experimentally but are only predicted to produce these proteins based on sequence similarity to other organisms.

| Locus | Sequence | Description | E-value | PI | Accession | Hit start | Hit end |
| --- | --- | --- | --- | --- | --- | --- | --- |
| 84800 | TGCAGCCAGACGTGCCCGCAGTGCGTGCACAGCAGCGGCCCCTGCCACCACATCACCGAGAT | PREDICTED: Taeniopygia guttata multiple EGF-like-domains 10 (MEGF10), mRNA | 3E-17 | 95 | XM_002188758.3 | 2369 | 2424 |
| 47743 | TGCAGGGCGCCCATGTCGTAGCGGTGCATGGGCTGCGGGTTGTGCGGGTGGTGCGGGTGATG | PREDICTED: Taeniopygia guttata SRY (sex determining region Y)-box 1 (SOX1), mRNA | 3E-23 | 98 | XM_030280051.2 | 1291 | 1352 |
| 111006 | TGCAGGTGGGTGATCTCTCCCAGCATGGTGAGCTCTCCGTTCTGAAGAGACAAGAGGCAGCA | PREDICTED: Taeniopygia guttata MTSS I-BAR domain containing 2 (MTSS2), mRNA | 2E-12 | 98 | XM_030282520.2 | 2115 | 2156 |
| 75123 | TGCAGGCATTTAAAGGACATGTAGATGTGGCACCTGGCGACATGGTTCAGTGGTGGCCTTGG | PREDICTED: Taeniopygia guttata uncharacterized LOC115494015 | 6E-13 | 89 | XR_003959071.2 | 11026 | 11080 |

**Table S3.** Results from STRUCTURE HARVESTER with analysis of all samples. When the highest LnP(*K*) (BOLD) is not *K* = 1, the highest Delta *K* is used to determine *K* (BOLD).

| *K* | Reps | Mean LnP(*K*) | Stdev LnP(*K*) | Delta *K* |
| --- | --- | --- | --- | --- |
| 1 | 3 | -800038.30 | 5.345 | - |
| 2 | 3 | -798361.90 | 8.141 | **147.695** |
| 3 | 3 | -797887.83 | 41.600 | 76.928 |
| 4 | 3 | -800613.97 | 2253.739 | 0.410 |
| 5 | 3 | -802417.10 | 2275.655 | - |

**Table S4.** The loading and model contribution of the percentage plant cover of eight dominant shrub species across three zones where TBGWs were present or absent. Factor scores were calculated using inverse transformed variables with varimax rotation. Shrub species with the highest factor loadings are shown in bold.

| Shrub species | PC1 | PC2 | PC3 | Communalities |
| --- | --- | --- | --- | --- |
| *Zygochloa paradoxa* | **-0.850** | 0.059 | 0.131 | 0.744 |
| *Atriplex vesicaria* | **0.677** | 0.141 | 0.084 | 0.486 |
| *Maireana aphylla* | 0.233 | **-0.799** | 0.136 | 0.711 |
| *M. astrotricha* | 0.434 | **0.786** | 0.066 | 0.810 |
| *M. pyramidata* | 0.458 | **0.597** | 0.401 | 0.727 |
| *Rhagodia spinescens* | 0.120 | -0.236 | **0.731** | 0.604 |
| *Acacia spp* | -0.295 | 0.014 | **0.726** | 0.614 |
| *A. nummularia omissa* | -0.105 | -0.225 | **-0.581** | 0.399 |

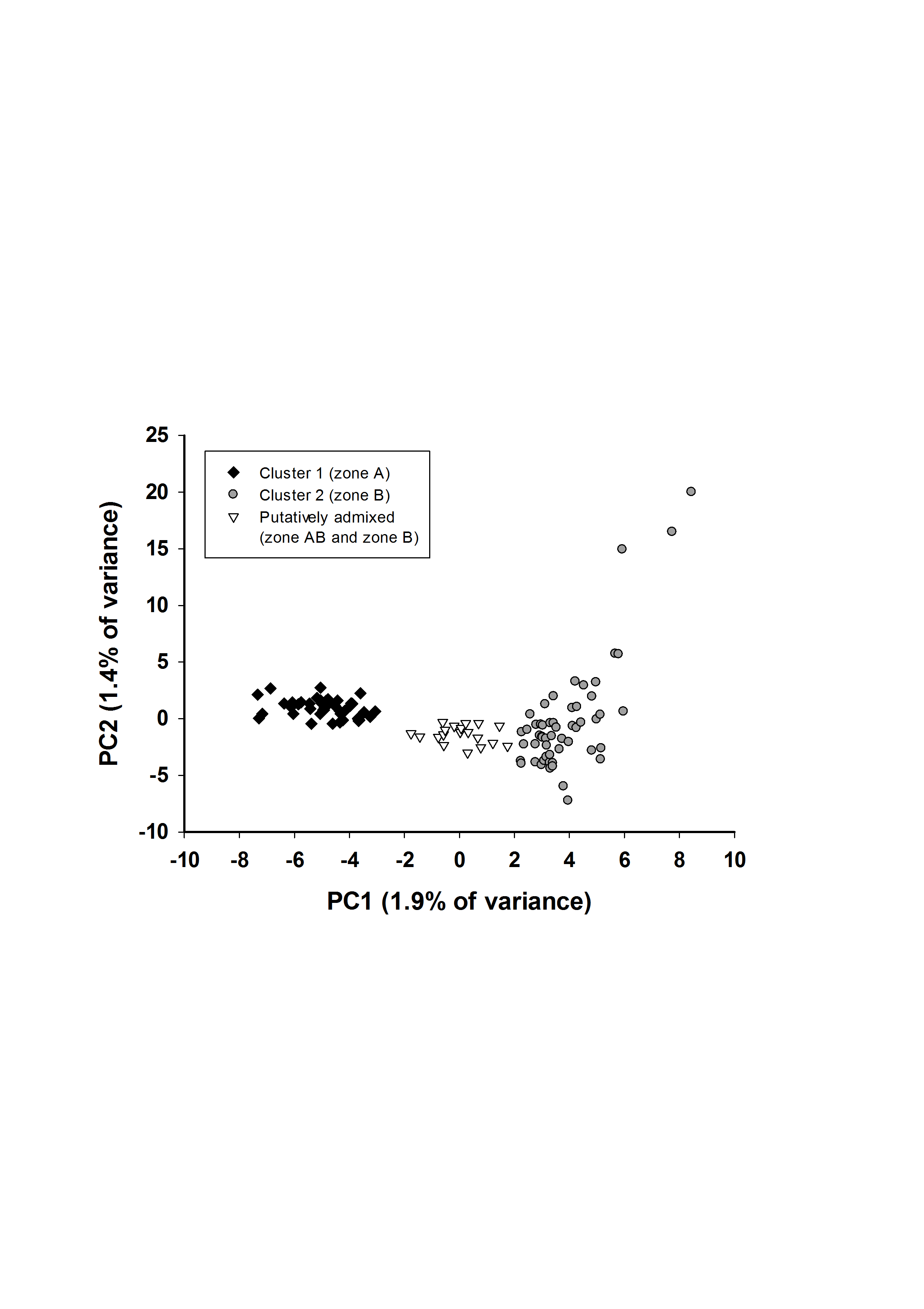

**Figure S1.** Plot of preliminary PCA with 13,635 loci used to identify putatively admixed individuals.

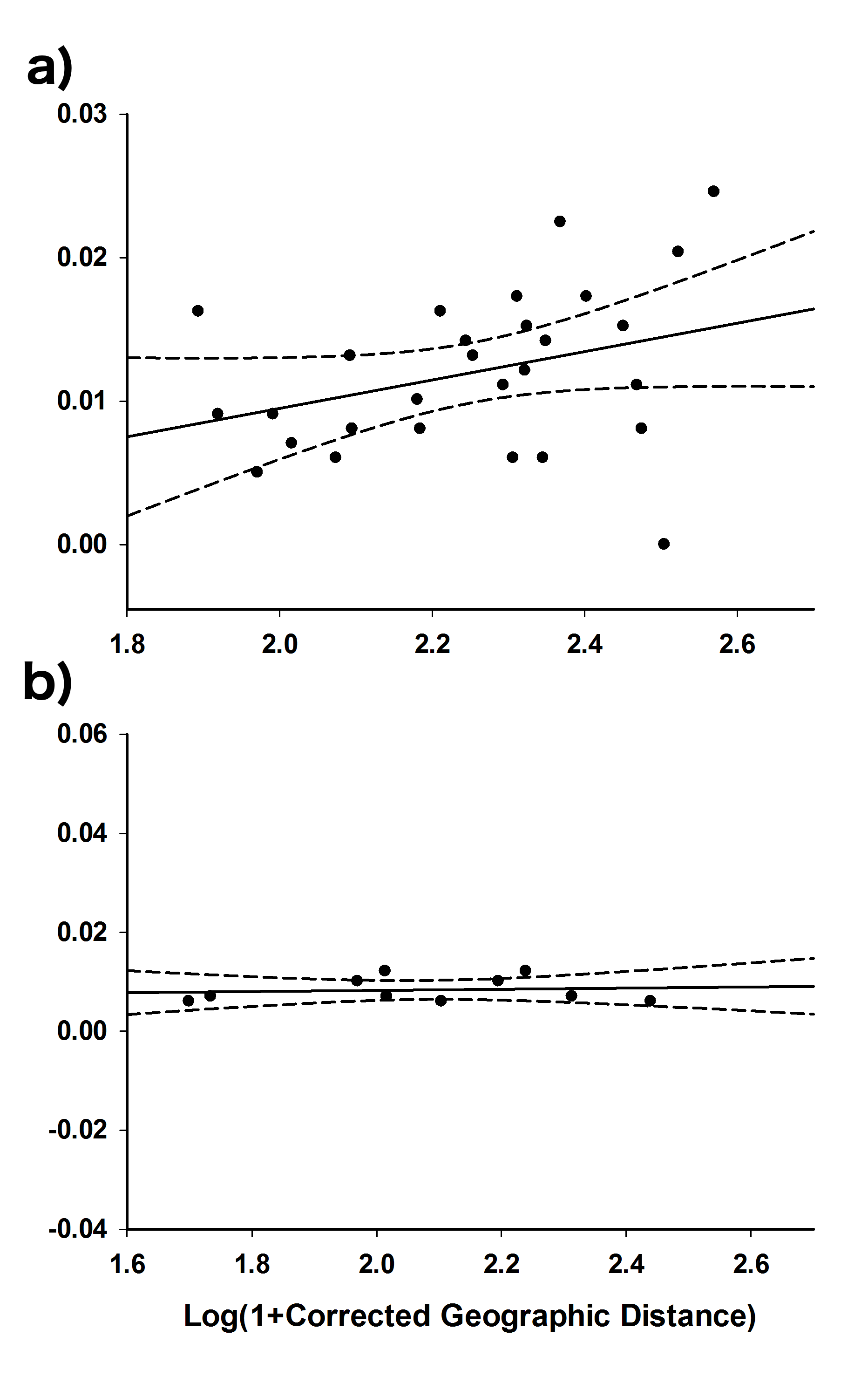

**Figure S2.** The pairwise genetic (*F*_ST_/(1 – *F*_ST_)) and geographic (log(1 + km)) relationship between localities by zone (zone A: *n* = 6, zone B: *n* = 3, and zone AB: *n* = 2) using a Mantel test. There was only one sample collected at the locality MTB, therefore this locality was excluded. The solid line is the line of best fit and the broken lines are the 95% confidence intervals. Different graphs represent comparisons of localities from different zones; a) zone A and zone AB, R^2^ = 0.112 and b) zone B and zone AB, R^2^ = 0.012.

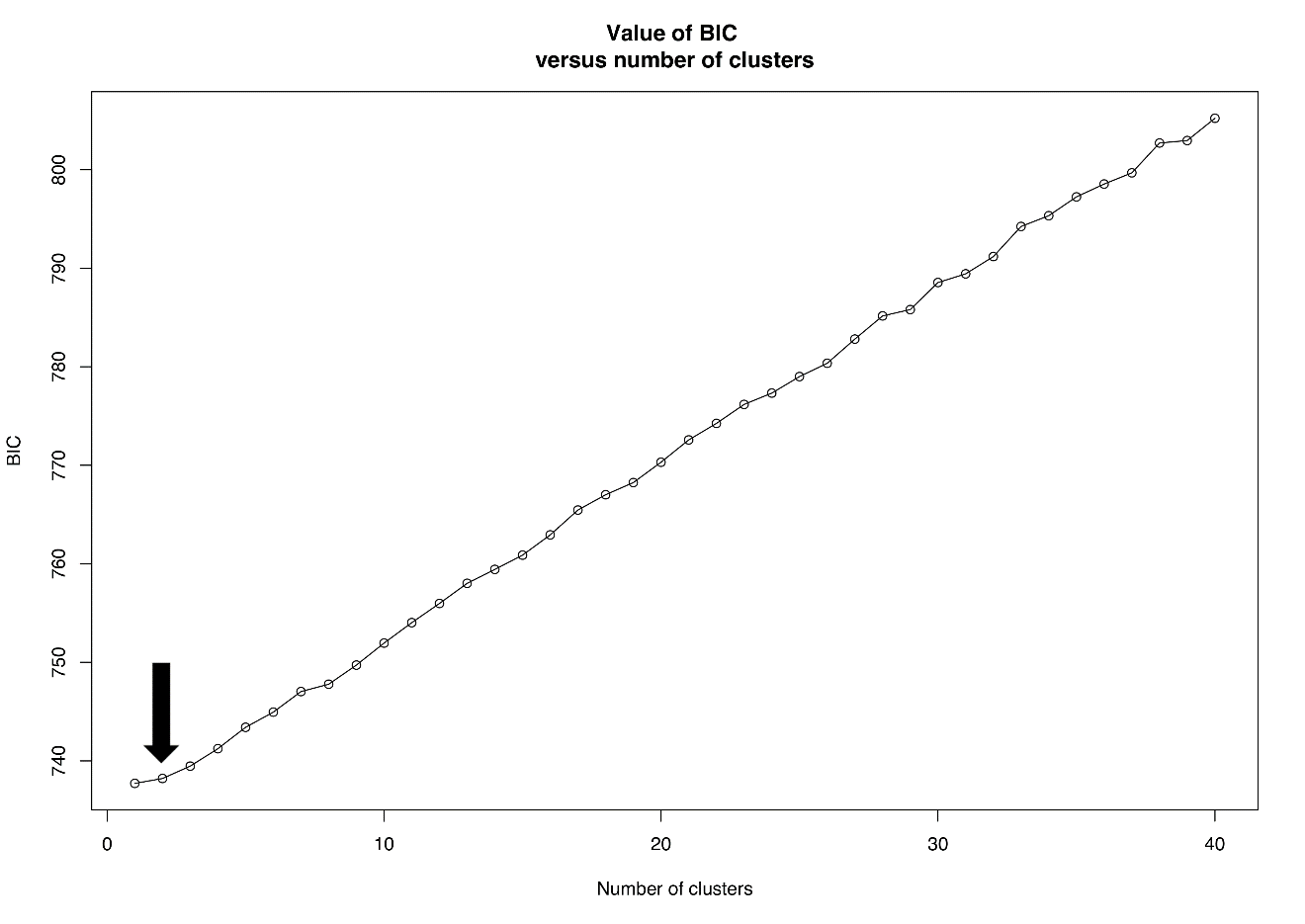
**Figure S3.** Results from the DAPC analysis. *K* is inferred from the lowest BIC value that falls after an elbow in the curve of BIC values as a function of *K* (arrow).

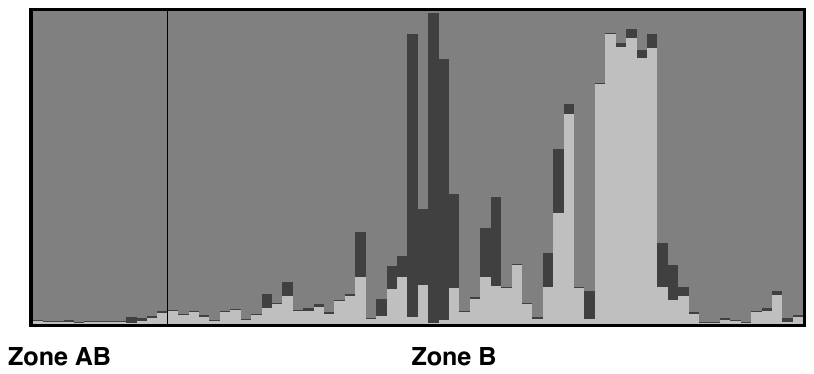

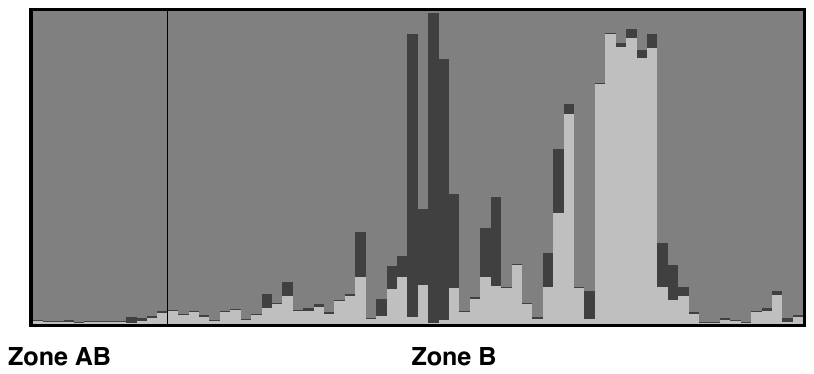

**Figure S4.** STRUCTURE results with only individuals from zone B and zone AB (*n* = 74). Delta *K* showed the most likely *K* = 3. The two small clusters within zone B are likely to belong to distantly related individuals. The smallest cluster of three individuals are also grouped separately on the PCA for PC2 (Figure 4).
